# Spatial inter-centromeric interactions facilitated the emergence of evolutionary new centromeres

**DOI:** 10.1101/2020.02.07.938175

**Authors:** Krishnendu Guin, Yao Chen, Radha Mishra, Siti Rawaidah B. M. Muzaki, Bhagya C. Thimmappa, Caoimhe E. O’Brien, Geraldine Butler, Amartya Sanyal, Kaustuv Sanyal

## Abstract

Centromeres of *Candida albicans* form on unique and different DNA sequences but a closely related species, *Candida tropicalis*, possesses homogenized inverted repeat (HIR)-associated centromeres. To investigate the mechanism of centromere-type transition, we improved the fragmented genome assembly and constructed a chromosome-level genome assembly of *C. tropicalis* by employing PacBio sequencing, chromosome conformation capture sequencing (3C-seq), chromoblot, and genetic analysis of engineered aneuploid strains. Further, we analyzed the 3D genome organization using 3C-seq data, which revealed spatial proximity among the centromeres as well as telomeres of seven chromosomes in *C. tropicalis*. Intriguingly, we observed evidence of inter-centromeric translocations in the common ancestor of *C. albicans* and *C. tropicalis*. Identification of putative centromeres in closely related *Candida sojae, Candida viswanathii* and *Candida parapsilosis* indicates loss of ancestral HIR-associated centromeres and establishment of evolutionary new centromeres (ENCs) in *C. albicans*. We propose that spatial proximity of the homologous centromere DNA sequences facilitated karyotype rearrangements and centromere type transitions in human pathogenic yeasts of the CUG-Ser1 clade.

## Introduction

The efficient maintenance of the genetic material and its propagation to subsequent generations determine the fitness of an organism. Genomic rearrangements are often associated with the development of multiple diseases, including cancer. Chromosomal rearrangements, on the other hand, are often observed during speciation (1). Such structural changes begin with the formation of at least one DNA double-strand break (DSB), which is generally repaired by homologous recombination (HR) or non-homologous end joining (NHEJ) *in vivo*. Studies using engineered *in vivo* model systems suggested that the success of DSB repair through HR depends upon an efficient identification of a template donor. This process of ‘homology search’ is facilitated by the physical proximity and the extent of DNA sequence homology (2-4). Multi-invasion-induced rearrangements (MIRs) involving more than one template donors have recently been shown to be influenced by physical proximity and homology (5). Therefore, the nature of genomic rearrangements is mostly dependent on the type of spatial genome organization. In yeasts, apicomplexans, and certain plants, centromeres cluster inside the nucleus (6), which may facilitate translocations between two chromosomes involving their centromeric and adjacent pericentromeric loci.

The centromere, one of the guardians of genome stability, assembles a large DNA-protein complex to form the kinetochore, which ensures fidelity of chromosome segregation by correctly attaching chromosomes to the spindle. Paradoxically, this conserved process of chromosome segregation is carried out by highly diverse species-specific centromere DNA sequences. For example, the length of centromere DNA is ∼125 bp in budding yeast *Saccharomyces cerevisiae* (7), but it can be as long as a few megabases in humans (8). Centromeres have been cloned and characterized from a large number of fungal species. The only factor that remains common to most fungal centromeres is the presence of histone H3 variant CENP-A^Cse4^ except in some Mucorales like *Mucor circinelloides* (9). Many kinetochore proteins are believed to have evolved from pre-eukaryotic lineages and remained conserved within closely related species complexes or expanded through gene duplication (10-12). It remains a paradox that despite the rapid evolution of centromere DNA, the kinetochore structure remains relatively well-conserved (13). Therefore, an examination of the evolutionary processes driving species-specific changes in centromere DNA is essential for a better understanding of centromere biology.

The first cloned centromere that of the budding yeast *S. cerevisiae* carries conserved genetic elements capable of forming a functional centromere *de novo* when cloned into a yeast replicative plasmid (7). Such genetic regulation of centromere function also exists in the fission yeast *Schizosaccharomyces pombe*, where centromeres possess inverted repeat-associated structures of 40-100 kb (14). Other closely related budding and fission yeasts were also found to harbor a DNA sequence-dependent regulation of centromere function (15-17), but the advantage of having such genetic regulation is not well understood. In fact, the majority of species with known centromeres are thought to be regulated by an epigenetic mechanism (13). A truly epigenetically-regulated fungal centromere carrying a 3-5 kb long CENP-A^Cse4^-bound unique DNA sequence exists in another budding yeast *C. albicans* (18), a CUG-Ser1 clade species in the fungal phylum of Ascomycota. Subsequently, such unique centromeres were also discovered in closely related *Candida dubliniensis* (19) and *Candida lusitaniae* (20). Strikingly, all seven centromeres of *C. tropicalis*, another CUG-Ser1 clade species, carry 3-4 kb long inverted repeats (IR) flanking ∼3 kb long CENP-A^Cse4^ rich central core (CC). The centromere sequences are highly identical to each other in *C. tropicalis*. Intriguingly, centromere DNA of *C. tropicalis* can facilitate *de novo* recruitment of CENP-A^Cse4^ to some extent (21). In contrast, centromeres of *C. albicans* completely lack such a DNA sequence-dependent mechanism (22). Such a rapid transition in the structural and functional properties of centromeres within two closely related species offers a unique opportunity to study the process of centromere type transition.

Kinetochore proteins appeared as a single punctum at the periphery of a nucleus indicating the presence of constitutively clustered centromeres in *C. tropicalis* (21). Our previous analysis also showed that centromeres of *C. tropicalis* were located near interchromosomal synteny breakpoints (ICSBs) as relics of ancient translocations in the common ancestor of *C. tropicalis* and *C. albicans* (21). Do homologous centromere DNA regions in close spatial proximity facilitate chromosomal translocation events? Due to the nature of the then-available fragmented genome assembly, the genome-wide distribution of the ICSBs and the spatial organization of the genome in *C. tropicalis* remained unexplored. However, the near-complete *C. albicans* genome assembly was available. Therefore, to examine whether the spatial proximity of clustered centromeres drives interchromosomal translocation events guiding speciation in the CUG-Ser1 clade required a chromosome-level complete genome assembly of *C. tropicalis*.

In this study, we constructed a chromosome-level gapless genome assembly of the *C. tropicalis* type strain MYA-3404 by combining information from previously available contigs, NGS reads and high-throughput 3C-seq data. Using this assembly and 3C-seq data, we studied the spatial genome organization in *C. tropicalis.* Next, we mapped the ICSBs in the *C. tropicalis* genome with reference to that of *C. albicans* (ASM18296v3) to test whether the frequency of ICSB correlated with the spatial genome organization. In addition, we performed Oxford Nanopore and Illumina sequencing and assembled the genome of *Candida sojae* (strain NCYC-2607), a sister species of *C. tropicalis* in the CUG-Ser1 clade (23). Finally, using this genome assembly of *C. sojae* and publicly available genome assembly of *C. viswanathii* (ASM332773v1), we identified the putative centromeres of these two species as HIR-associated loci syntenic to the centromeres of *C. tropicalis*. Based on our results, we propose a model that suggests homology and proximity guided centromere-proximal translocations facilitated karyotype evolution and possibly aided in rapid transition from HIR-associated to unique centromere types in the members of the CUG-Ser1 clade.

## Results

### A chromosome-level gapless assembly of the *C. tropicalis* genome in seven chromosomes

*C. tropicalis* has seven pairs of chromosomes (21, 24). However, the current publicly available genome assembly (ASM633v3) has 23 nuclear contigs and one mitochondrial contig. To completely assemble the nuclear genome of *C. tropicalis* in seven chromosomes, we combined results of short-read Illumina sequencing and long-read single molecule real-time sequencing (SMRT-seq) with high-throughput 3C-seq (simplified Hi-C) experiment (Figure 1A, S1A-D) (25). We started from the publicly available genome assembly of *C. tropicalis* strain MYA-3404 in 23 nuclear contigs (ASM633v3, Assembly A) (24). We used Illumina sequencing reads to scaffold them into 16 contigs to get Assembly B (Figure 1A). Next, we used the SMRT-seq long reads to join these contigs, which resulted in an assembly of 12 contigs (Assembly C, Table S1). Based on the contour clamped homogenized electric field (CHEF)-gel karyotyping (Figure 1B) and 3C-seq data (Figure S1E-G), we joined two contigs and rectified a misjoin in Assembly C to produce an assembly of seven chromosomes and five short orphan haplotigs (OHs). We suspected that the OHs are heterozygous loci in the diploid genome of *C. tropicalis*. Analysis of the *de novo* contigs (Figure S1H, Methods), sequence coverage data (Figure S2A-B), and Southern hybridization of engineered aneuploid strains demonstrated that the small OHs mapped to heterozygous regions of the genome (Figure S2C-I, Methods). Next, we used *de novo* contigs to fill pre-existing 104 N-gaps and scaffolded 14 sub-telomeres (Figure S3A-C, Table S2). Finally, we used 3C-seq reads to polish the complete genome assembly of *C. tropicalis* constituting 14,609,527 bp in seven elomere-to-telomere long gapless chromosomes (Figure 1B). We call this new assembly as Assembly2020.

**Figure 1.**
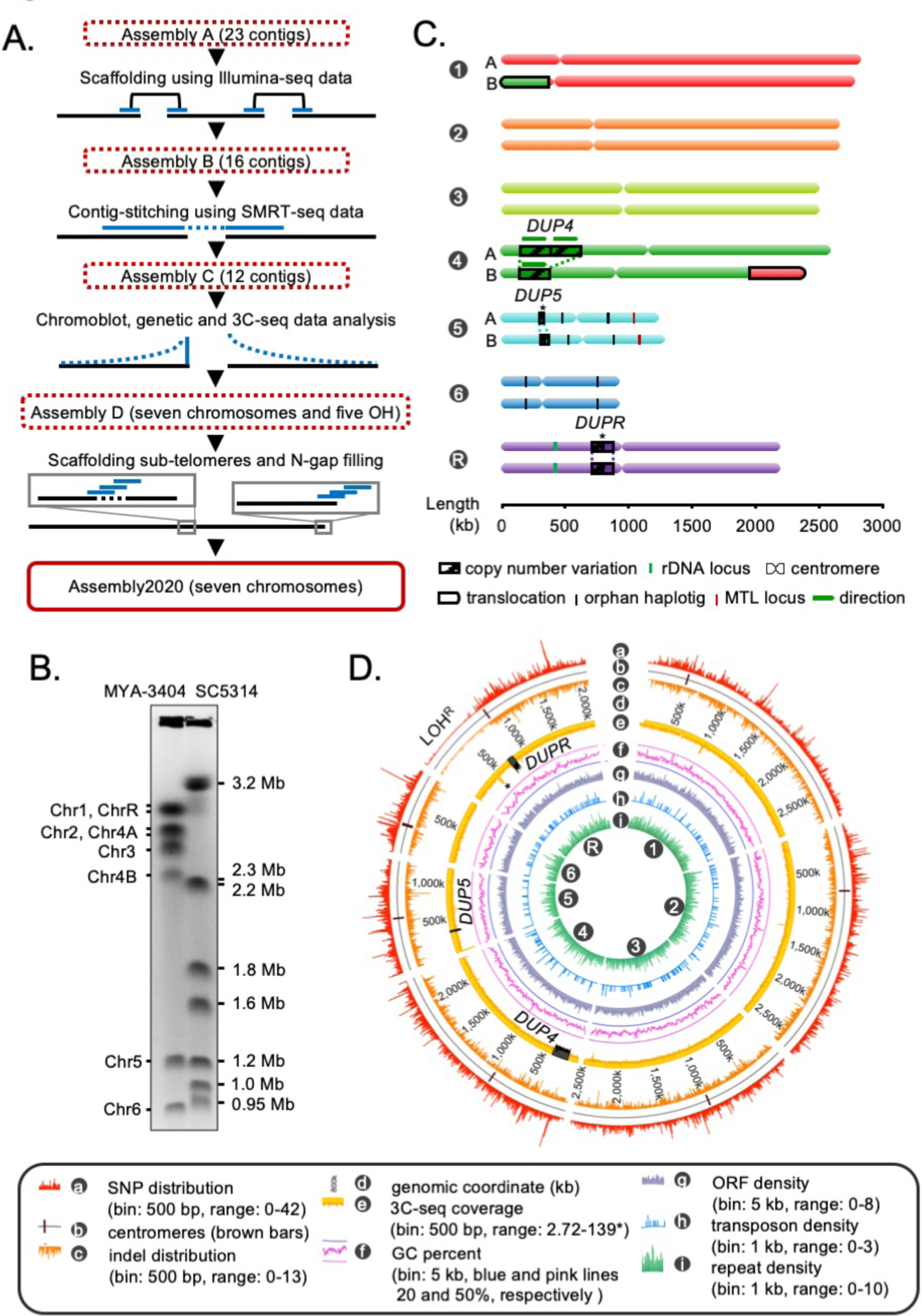
Construction of the gapless assembly of *C. tropicalis* type strain MYA-3404 in seven chromosomes. A. Schematic showing the stepwise construction of the gapless chromosome-level assembly (Assembly2020) of *C. tropicalis* (also see Figure S1 and S2). B. An ethidium bromide (EtBr)-stained CHEF gel image of separated chromosomes of the *C. tropicalis* (strain MYA-3404) and *C. albicans* (strain SC5314) (Methods). *C. albicans* chromosomes are used as size markers for estimation and validation of lengths and identities of *C. tropicalis* chromosomes in the newly constructed Assembly2020. C. An ideogram of seven chromosomes of *C. tropicalis* as deduced from Assembly2020 and drawn to scale. The genomic location of the three loci showing copy number variations (CNVs), *DUP4, DUP5* and *DUPR* located on Chr4, Chr5 and ChrR respectively, are marked and depicted as striped box. The CNVs for which the correct homolog-wise distribution of the duplicated copy is unknown are marked with asterisks. Homolog-specific differences for Chr1 and Chr4, occurred due to an exchange of chromosomal parts in a balanced heterozygous translocation between Chr1B and Chr4B, are highlighted with black borders (also see Figure S4C). D. A circos plot showing the genome-wide distribution of various sequence features. Very high sequence coverage at rDNA locus is clipped for more precise representation and marked with an asterisk.

We assigned the numbers to each chromosome according to the length, starting from the longest as chromosome 1 (Chr1) through the shortest as chromosome 6 (Chr6). The remaining chromosome, the one containing the rDNA locus, was named as chromosome R (ChrR) (Figure 1C). Accordingly, centromeres on each chromosome were named after the respective chromosome number. Additionally, we oriented the DNA sequence of each chromosome in a way to consistently maintain the short arm at the 5′ end. The statistics of these genome assemblies of *C. tropicalis* is summarized in Table S3. In Assembly2020, 1278 out of 1315 Ascomycota-specific BUSCO gene sets could be identified compared to 1255 identified using Assembly A (Table S4, Methods). The inclusion of 23 additional BUSCO gene sets suggests significantly improved contiguity and completeness of Assembly2020.

Previously, using centromere-proximal probes, we could distinctly identify five chromosomes (Chr1, Chr2, Chr3, Chr5, and Chr6) in chromoblot analysis (21). However, the lengths of Chr4 and ChrR could not be determined. To validate the correct assembly of these two chromosomes (Chr4 and ChrR), we performed additional chromoblot analysis. We observed that Chr4 homologs differed in size (Figure S4A). Analysis of the sequence coverage across Chr4 identified a substantial internal duplication of ∼235 kb region, which could explain the size difference between the homologs Chr4A and Chr4B (Figure 1C, S4B). We named this duplicated locus as *DUP4.* Subsequently, we scanned the entire genome for the presence of copy number variations (CNVs), which led to the identification of two additional large-scale duplication events: one each on Chr5 (*DUP5*, ∼23 kb) and ChrR (*DUPR*, ∼80 kb) (Figure 1C, S4B). Further, using CNAtra software (26) we confirmed these duplication events and identified additional small-scale CNV loci with copy number <1.5 or >2.5 (Figure S4C). Additionally, we detected a balanced heterozygous translocation event between Chr1 and Chr4 (Figure S5A) through analyses of 3C-seq data and *de novo* contigs (Figure S5B). This translocation was validated using chromoblot analysis (Figure S5C) as well as Illumina, and SMRT-seq read mapping (Figure S5D). Thus, while chromoblot analysis suggests that the actual length of ChrR is ∼2.8 Mb (Figure S5E), the assembled length is 2.1 Mb (Figure 1C). Considering the length of the rDNA locus is ∼700 kb in *C. albicans* (27), we reason that the difference between the assembled length and actual length (derived from chromoblot analysis) of ChrR in *C. tropicalis* can be attributed to the presence of the repetitive rDNA locus of ∼700 kb, which is not completely assembled in Assembly2020.

Next, we performed phasing of the diploid genome of *C. tropicalis* using SMRT-seq and 3C-seq data to identify the homolog-specific variations (Methods). This analysis produced 16 nuclear contigs, which were colinear with the chromosomes of Assembly2020, except for the previously validated heterozygous translocation between Chr1 and Chr4 (Figure S5F). To characterize the sequence variations in the diploid genome of *C. tropicalis*, we identified the single nucleotide polymorphisms (SNPs) and insertion-deletion (indel) mutations (Methods). Intriguingly, we detected a long chromosomal region depleted of SNPs and indels on the left arm of ChrR (Figure 1D). We named this region that lost heterozygosity on ChrR as LOH^R^. Strikingly, we found parts of the syntenic region of LOH^R^ to be SNP and indel depleted in the *C. sojae* strain NCYC-2607, a closely related species of *C. tropicalis*, as well as in *C. albicans* reference strain SC5314 (Figure S6). We also identified the genome-wide distribution of transposons and simple repeats but could not detect preferential enrichment of these sequence elements at any specific genomic location in *C. tropicalis* (Figure 1D). Together, we demonstrate, for the first time, multiple CNVs, a long-track LOH, and evidence of a heterozygous reciprocal translocation event in the diploid genome of *C. tropicalis.* Possible implications of these events in conferring virulence and drug resistance in this successful human fungal pathogen remain to be explored.

### Conserved principle of the spatial genome organization in *C. tropicalis* and *C. albicans*

Indirect immunofluorescence imaging of the *C. tropicalis* strain (CtKS102) expressing Protein-A tagged CENP-A^Cse4^ suggested that centromeres are clustered and localized at the periphery of the DAPI-stained nuclear DNA mass as a single punctum (Figure 2A-B). We mapped 3C-seq data (see Methods), that were generated using DpnII, to the Assembly2020 to construct the genome-wide chromatin contact map of *C. tropicalis*. The resultant heatmap depicts high signal intensities along the diagonal, indicating that the intrachromosomal interactions are generally stronger than interchromosomal interactions, as observed before (Figure 2C) (28). However, the most striking feature of the heatmap is the presence of conspicuous puncta in the interchromosomal areas, which signify strong spatial proximity between centromeres (Figure 2C-D). The aggregate signal analysis further reiterated the enrichment of centromere-centromere interactions (Figure 2E). Strikingly, we also noted the enrichment of telomere-telomere interactions as compared to the neighboring regions (Figure 2C-E). Statistical comparison was then performed between these telomere-telomere interactions and bulk chromatin, which revealed that the interchromosomal telomeric interactions were significantly greater than the all interchromosomal interactions (Mann-Whitney U test P value = 1.129.10^−11^) (Figure S7A). On the other hand, *cis* interactions between the two telomeres of an individual chromosome (intrachromosomal telomeric interactions) were also significantly enhanced compared to all intrachromosomal long-range (>100 kb) interactions (Mann-Whitney U test P value = 7.374·10^−11^) (Figure S7B). All these lines of evidence prompted us to propose that *C. tropicalis* chromosomes adopt the Rabl-like configuration, a characteristic feature of the higher-order genome organization in yeasts (28-30).

**Figure 2.**
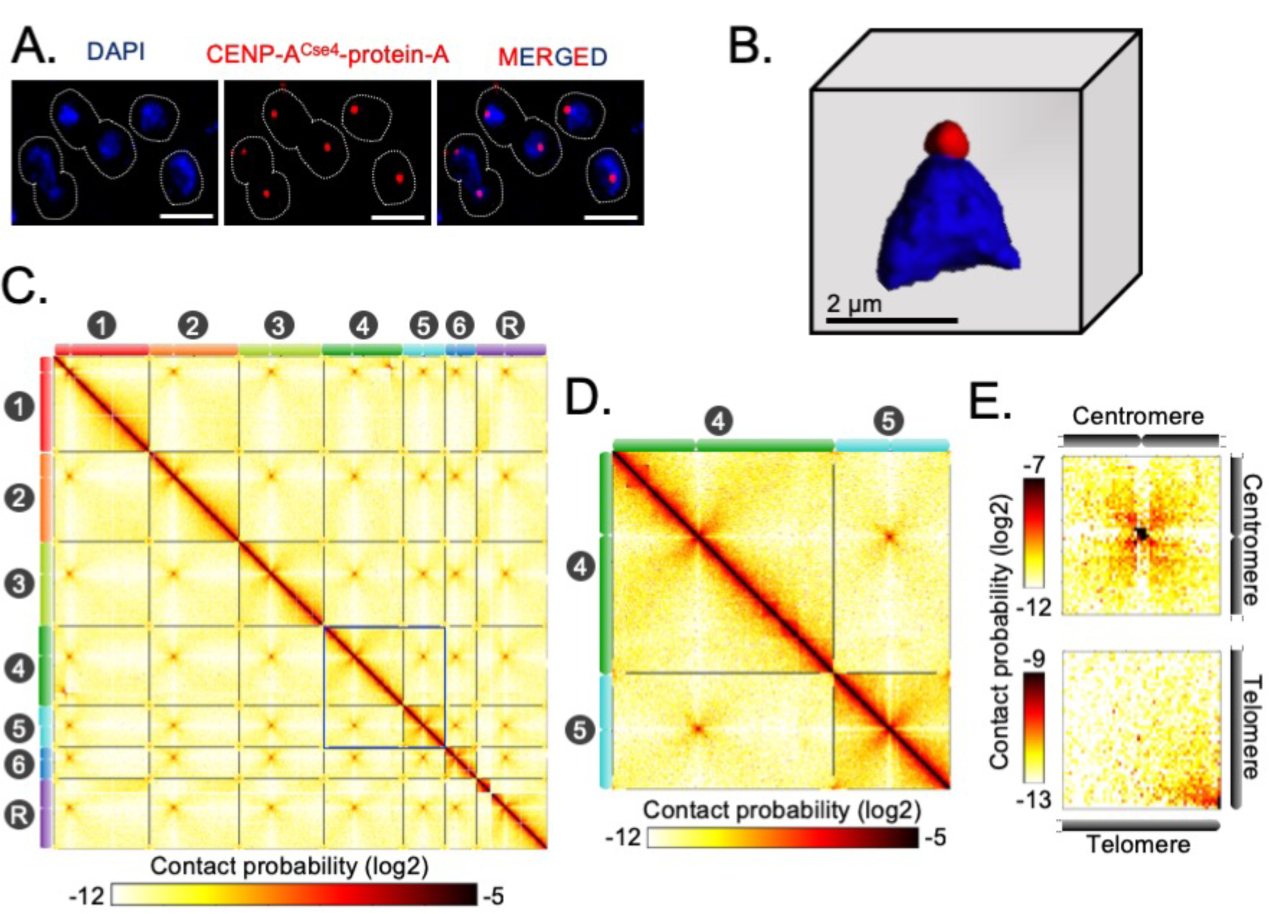
Spatial genome organization reveals centromere-centromere and telomere-telomere contacts in *C. tropicalis.* A. A representative field image of *C. tropicalis* (strain CtKS102) cells expressing Protein-A tagged CENP-A^Cse4^. CENP-A signals (blue) were obtained using anti-Protein A antibodies by indirect immuno-fluorescence microscopy. Nuclei of the corresponding cells were stained by DAPI (blue). The images were acquired using a DeltaVision imaging system (GE) and processed using FIJI software (93). Scale, 2 μm. B. A 3D reconstruction showing clustered kinetochores marked by CENP-A^Cse4^ (red) at the periphery of the DAPI-stained nucleus (blue) using Imaris software (Oxford Instruments) in *C. tropicalis*. Scale, 2 μm. C. A genome-wide contact probability heatmap (bin size = 10 kb) generated using 3C-seq data. Chromosome labels and their corresponding ideograms are shown on the axes of the heatmap. Colorbar represents the contact probability in the log2 scale. D. Zoom in view of heatmap showing Chr4 and Chr5 from panel C (blue box). E. Heatmaps plotted from aggregate signal analysis of matrices (bin size = 2 kb) surrounding centromere-centromere (top) or telomere-telomere interactions (bottom). *Top*, genomic loci containing mid-points of centromeres are aligned at the center (red bar); *bottom*, genomic loci from 5′ or 3′ ends of chromosomes are aligned at the bottom right corner.

Previously, microscopic and Hi-C studies revealed similar centromere clustering and strong physical interactions among centromeres in *C. albicans* (30-32). This study now reveals that despite substantial karyotypic changes, a conserved principle of genome organization exists in two yeast species, *C. albicans* and *C. tropicalis*, with diverged centromere features.

### Centromere and telomere proximal loci are hotspots for complex translocations

Using the chromosome-level assemblies of *C. tropicalis* type strain MYA-3404 and *C. albicans* type strain SC5314 (ASM18296v3), we performed a detailed genome-wide synteny analysis employing four different approaches. We used two analytical tools, Symap (33) and Satsuma synteny (34), and a custom approach to identify the ICSBs based on the synteny of the conserved orthologs (Figure 3A). Next, we compared and validated the results obtained from our custom approach of analysis with another published tool Synchro (35). Considering the *C. albicans* genome as the reference, all four methods of analyses suggest that six out of seven centromeres (except *CEN6*) of *C. tropicalis* are located proximal to multiple ICSBs (Figure 3A, S8A). Although it appears that *CtCEN6* escaped inter-centromeric translocations, synteny analysis suggested that a chromosomal region carrying three consecutive *CtCEN6*-proximal ORFs was lost in the *C. albicans* genome (Figure S8B). Strikingly, these ICSBs are rare at the chromosomal arms (Figure 3A). ORF-level synteny analysis further revealed that four out of seven centromeres (*CEN2, CEN3, CEN5*, and *CENR*) in *C. tropicalis* are precisely located at the ICSBs (Figure S8C), while multiple ICSBs are located within ∼100 kb of other two centromeres (Figure 3A). Additionally, a convergence of orthoblocks from as many as four different chromosomes of *C. albicans* was detected within 100 kb of *C. tropicalis* centromeres (Figure 3B). It is important to note that by using the *C. tropicalis* genome as the reference, all centromeres of *C. albicans*, except *CaCEN2*, were found to be associated with ICSBs (Figure S8D). Taken together, centromeres of both these species are found to be associated with chromosomal translocations.

**Figure 3.**
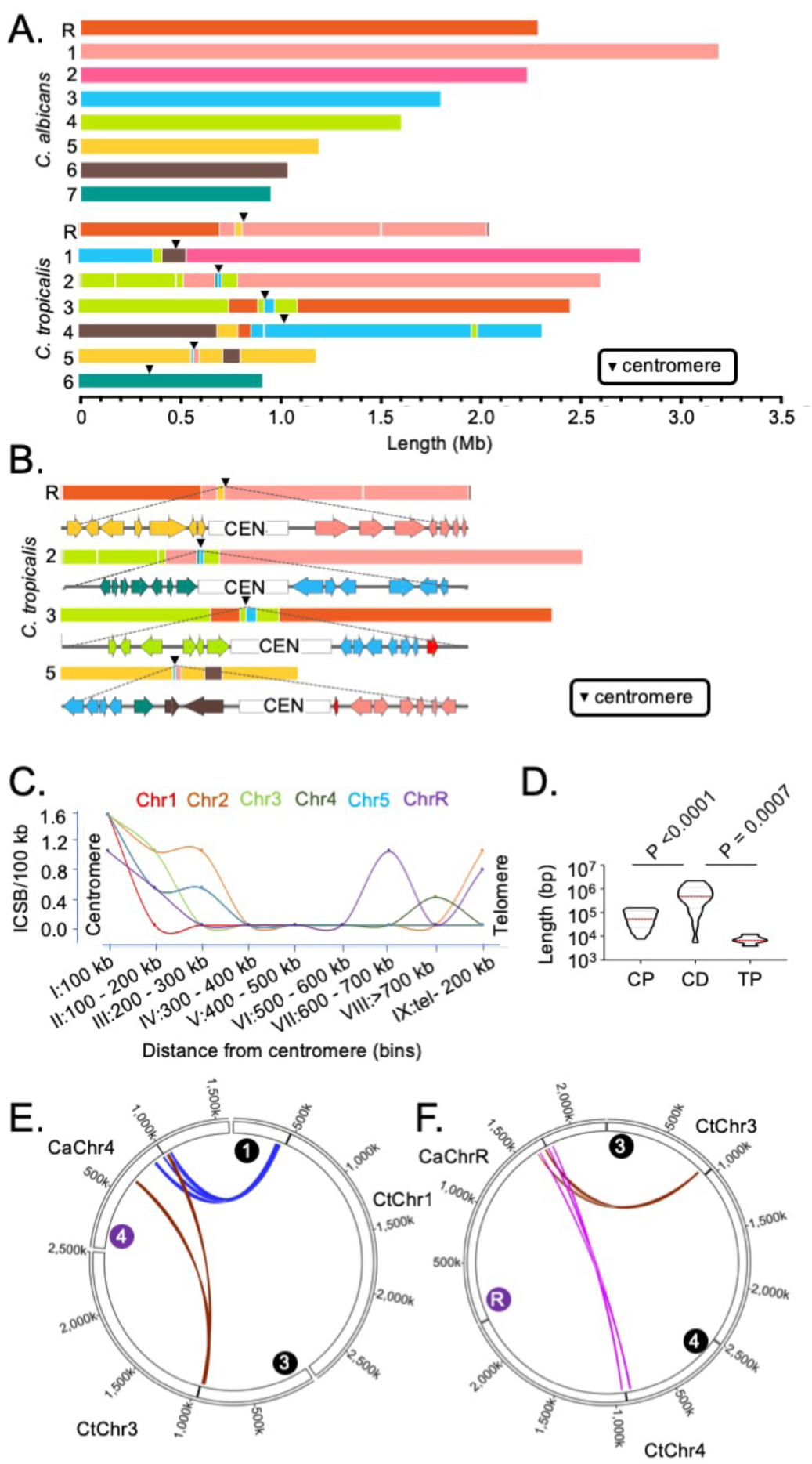
Genome-wide mapping of interchromosomal synteny breakpoints in *C. tropicalis* identifies a spatial cue for karyotype evolution. A. Scaled representation of the color-coded orthoblocks (relative to *C. albicans* chromosomes) and ICSBs (white lines) in *C. tropicalis* (Methods). Orthoblocks are defined as stretches of the target genome (*C. tropicalis*) carrying more than two syntenic ORFs from the same chromosome of the reference genome (*C. albicans*). The centromeres are represented with black arrowheads. B. Zoom in view of the *C. tropicalis* centromere-specific ICSBs on *CEN2, CEN3, CEN5* and *CENR* showing the color-coded (relative to *C. albicans* chromosomes) ORFs flanking each centromere. *tropicalis-*specific unique ORFs proximal to *CEN3* and *CEN5* are shown in red. C. A plot showing the chromosome-wise ICSB density, calculated as number of ICSBs per 100 kb of the C. *tropicalis* genome (*y-*axis), as a function of the linear distance from the centromere in nine bins. These bins are a) 0 - 100 kb on both sides of centromere (bin I), b) 100 - 200 kb (bin II), c) 200 - 300 kb (bin III), d) 300 - 400 kb (bin IV), e) 400 - 500 kb (bin V), f) 500 - 600 kb (bin VI), g) 600 - 700 kb (bin VII), h) >700 kb to 200 kb from telomere ends (bin VIII), and i) 200 kb from the telomere ends (bin IX). Chr6 was excluded from this analysis, as it does not harbor any ICSB. A violin plot comparing the distribution of lengths of orthoblocks (*y-*axis) at three different genomic zones: a) the centromere-proximal zone (CP), b) the centromere-distal zone (CD), and c) telomere-proximal zone (TP). Orthoblocks, which span over more than one zone, were assigned to the zone with maximum overlap. The centromere-distal dataset was compared with the other two groups using the Mann-Whitney U test and the respective *P* values are mentioned. E - F. Circos plots representing the convergence of centromere-proximal ORFs of *C. tropicalis* chromosomes near the centromeres (*CEN4* and *CEN7*) of *C. albicans*. Chromosomes of *C. tropicalis* and *C. albicans* are marked with black and purple filled circles at the beginning of each chromosome, respectively.

To correlate the frequency of translocations with the spatial genome organization, we quantified ICSB density (the number of ICSBs per 100 kb of the genome) for different zones across the chromosome for all chromosomes except CtChr6 (Figure 3C). Our analysis reveals that the ICSB density is maximum at the centromere-proximal zones for all six chromosomes, but drops sharply at the chromosomal arms. However, the ICSB density near the telomere-proximal zone for Chr2, Chr4, and ChrR shows an increase compared to the chromosomal arms, albeit at a lower magnitude than centromeres. We also compared the lengths of orthoblocks across three different genomic zones - the centromere-proximal (0 - 300 kb from the centromere on both sides), centromere-distal (>300 kb from the centromere to 200 kb away from the telomere ends), and telomere-proximal (0 - 200 kb from the telomere ends) zones. This analysis further reveals that the lengths of the orthoblocks located proximal to centromeres and telomeres are significantly smaller than orthoblocks located at the centromere-/telomere-distal zones (Figure 3D).

We further probed into the consequences of strong inter-centromeric interactions, as described above. Synteny analysis across centromere-proximal regions of the two species hints that inter-centromeric translocations may have occurred in the common ancestor of *C. albicans* and *C. tropicalis*. If such is the case, the centromere-proximal ORFs of different chromosomes in *C. tropicalis* should have converged on the *C. albicans* genome. Indeed, we identified at least ten loci where a convergence of *C. tropicalis* ORFs from different chromosomes had taken place in *C. albicans* (Figure S8E). Intriguingly, we found four such loci that are proximal to the centromeres (*CEN3, CEN4, CEN7*, and *CENR*) in *C. albicans* (Figure 3E-F, S8F-G). This observation strongly supports the possibility of inter-centromeric translocation events in the common ancestor of *C. albicans* and *C. tropicalis.* Additionally, the other four centromeres in *C. albicans* are located proximal to ORFs, orthologs of which are also proximal to the centromeres in *C. tropicalis* (Figure S8E). We posit that the ancestral HIR-associated centromeres were lost in *C. albicans*, and ENCs formed proximal to the ancestral centromere loci on unique DNA sequences. A similar centromere type transition within two isolates of *C. parapsilosis*, another species of the CUG-Ser1 clade, has been recently reported (36).

### Rapid transition in the centromere type within the members of the CUG-Ser1 clade

Since multiple translocation events near centromeric regions of the *C. tropicalis* genome could be detected, we hypothesized that complex translocations between HIR-associated centromeres in the common ancestor of *C. albicans* and *C. tropicalis* led to the loss of HIR and the evolution of unique centromere types observed in *C. albicans* and *C. dubliniensis.* However, the genomic rearrangements are rare events, even at the evolutionary time scale. Therefore, if HIR-associated centromeres are to be the ancestral state from which unique centromeres were derived, some other closely related species should have retained HIR-associated centromeres. Indeed, we identified eight HIR-associated structures (Table S7), in the reference genome of *C. parapsilosis* strain CDC317 (ASM18276v2). Identification of the HIR-associated structures present at the intergenic and transcription-poor regions, one each on all eight chromosomes, suggests that these loci are the putative centromeres of *C. parapsilosis*. Indeed, it was recently reported that all eight CENP-A^Cse4^ enriched centromeres in the CLIB214 strain of *C. parapsilosis* are located at HIR-associated loci (36). Based on these lines of evidence, we conclude that the common ancestor of *C. albicans* and *C. tropicalis* possibly carried HIR-associated centromeres. Surprisingly, two centromeres in another isolate (90-137) of *C. parapsilosis* have been shown to be formed on non-HIR-associated loci (36). However, the driving force triggering polymorphisms in centromere locations within the same species is yet to be understood.

Although IRs are present in *CEN4, CEN5*, and *CENR* of *C. albicans*, these sequences are not homogenized like the HIR-associated centromeres in *C. tropicalis* (Figure 4A). To study the presence of HIRs in *C. sojae* (NCYC-2607), a sister species of *C. tropicalis* (23), we assembled its genome into 42 contigs, including seven chromosome-length contigs (Methods). Using this assembly, we identified seven putative centromeres in *C. sojae* as intergenic and HIR-associated loci syntenic to the centromeres in *C. tropicalis* (Figure S9A-B, D). Each of these seven putative centromeres in *C. sojae* consists of a ∼2 kb long CC region flanked by 3-12 kb long inverted repeats (Table S5). Using a similar approach, we identified six HIR-associated centromeres in the publicly available genome assembly (ASM332773v1) of *Candida viswanathii*, another species closely related to *C. tropicalis* (Figure S9C, E, Table S6) (37). A dot-plot analysis identified the presence of homologous sequences shared across IRs but not among the CC elements (Figure 4A) of the HIR-associated centromeres present in *C. tropicalis* and the putative centromeres of *C. sojae* and *C. viswanathii* (Table S7). Moreover, we detected extensive structural conservation in centromere DNA elements, especially among IRs within an individual species (Figure S10A). These structural feature of IRs are also significantly conserved across the three species, *C. tropicalis, C. sojae*, and *C. viswanathii* (Figure S10B).

**Figure 4.**
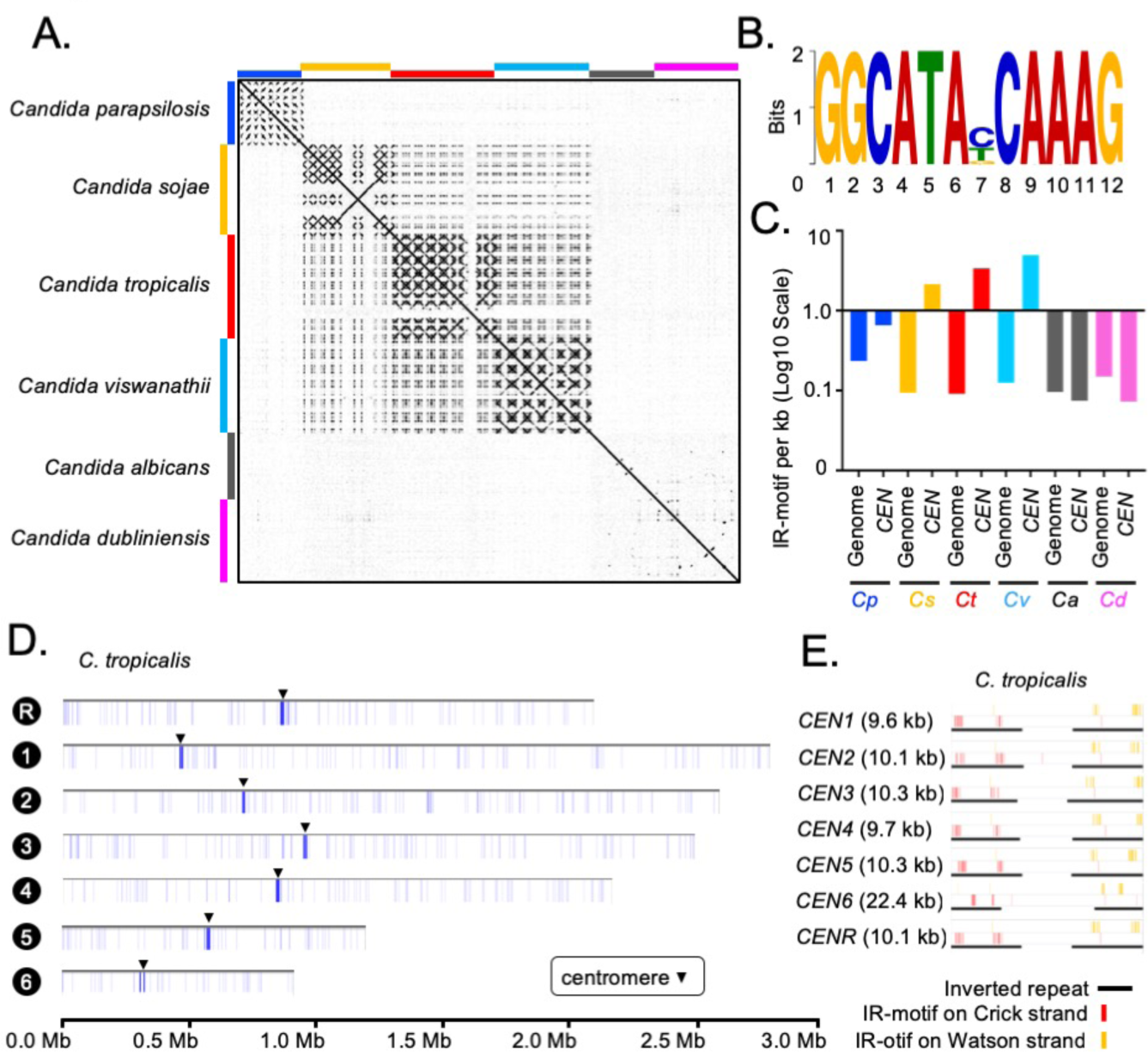
Genome-wide analysis of centromere DNA sequences across the CUG-Ser1 clade reveals the emergence of unique centromeres from an ancestral homogenized inverted repeat-associated centromere type. A. A dot-plot matrix representing the sequence and structural homology among species of the CUG-Ser1 clade was generated using Gepard (Methods). B. A logo plot showing the 12 bp-long IR-motif, identified using MEME-suit (Methods). C. The distribution of IR-motif density on centromere DNA sequences and across the entire genome of each species was calculated as the number of motifs per kb of DNA (Methods). Note that *C. albicans* and *C. dubliniensis* centromeres that form on unique and different DNA sequences do not contain the IR-motif. D. IGV track images showing the IR-motif density across seven chromosomes of *C. tropicalis.* The location of the centromere on each chromosome is marked with a black arrowhead. E. IGV track images showing the IR-motif distribution across seven HIR-associated centromeres of *C. tropicalis.*

Cloning of a full-length centromere of *C. tropicalis* in a replicative plasmid facilitated *de novo* CENP-A^Cse4^ deposition but failed to do so when the native IRs were replaced with Ca*CEN5* IRs (21). This result indicated DNA sequence specificity is required for centromere function in *C. tropicalis.* To identify the DNA sequence as a putative genetic element, we analyzed centromere DNA sequences of all three *Candida* species with HIR-associated centromeres and the unique centromeres of *C. albicans* for the presence of any conserved motif(s) (Methods). This analysis identified a highly conserved 12-bp motif (dubbed as IR-motif) (Figure 4B) clustered specifically at centromeres but not anywhere else in the entire genome of *C. tropicalis, C. sojae* and *C. viswanathii* (Figure 4C-D, S10C). On the contrary, the IR-motif density at centromeres in *C. albicans* remains approximately an order of magnitude lower than that of *C. tropicalis* (Figure 4C). This observation indicates a potential function of IR-motifs in the regulation of *de novo* CENP-A^Cse4^ loading in *C. tropicalis*. Moreover, this *CEN*-enriched motif found at IRs is absent at central core region in *C. tropicalis* (Figure 4E) and at the putative centromeres in *C. sojae* and *C. viswanathii* (Figure S10D). Additionally, we noted that the direction of the IR-motif is diverging away from the central core in *C. tropicalis* (Figure S10E) as well as in the other two species (Figure S10F). The conserved structure and organization of the IR-motif sequences in the HIR-associated centromeres of three *Candida* species suggest an inter-species conserved function of the IR DNA sequence. However, the clusters of IR-motifs are located at a variable distance from CC in these species (Figure S10G). The importance of the sequence and the density of IR-motifs on the centromere function is yet to be determined.

## Discussion

In this study, we improved the current genome assembly of the human fungal pathogen *C. tropicalis* by employing SMRT-seq, 3C-seq, and chromoblot experiments, and present Assembly2020, the first chromosome-level gapless genome assembly of this organism. We further identified three large-scale duplication events and few small-scale CNV loci in its genome, phased the diploid genome of *C. tropicalis*, and mapped SNPs and indels. We constructed a genome-wide chromatin contact map and identified significant centromere-centromere as well as telomere-telomere spatial interactions. Comparative genome analysis between *C. albicans* and *C. tropicalis* reveals that six out of seven centromeres of *C. tropicalis* are mapped precisely at or proximal to ICSBs. Strikingly, ORFs proximal to the centromeres of *C. tropicalis* are converged into specific regions on the *C. albicans* genome, suggesting that inter-centromeric translocations may have occurred in their common ancestor. Moreover, the presence of HIR-associated putative centromeres in *C. sojae* and *C. viswanathii*, like in *C. tropicalis*, suggests that such a centromere structure is plausibly the ancestral form in the CUG-Ser1 clade but lost both in *C. albicans* and *C. dubliniensis.* We propose that loss of such a centromere structure might have occurred during translocation events involving centromeres of homologous DNA sequences in the common ancestor, to give rise to ENCs on unique DNA sequences and facilitated speciation.

Unlike other centromeres, *CEN6* of *C. tropicalis* did not seem to undergo inter-centromeric translocations. A closer analysis revealed that three *CEN6*-associated ORFs of *C. tropicalis* are absent in the *C. albicans* genome while the other flanking ORFs remain conserved. This observation can be explained by a double-stranded DNA break at the centromere followed by the fusion of broken ends resulting in the loss of those ORFs.

The availability of the chromosome-level genome assembly and improved annotations of genomic variants and genes absent in the publicly available fragmented genome assembly of *C. tropicalis* should greatly facilitate genome-wide association studies to understand the pathobiology of this organism including the cause of antifungal drug resistance. Besides, this study sheds light on how genetic elements required for *de novo* centromere establishment in an ancestral species could be lost in the derived lineages to give rise to epigenetically-regulated centromeres.

*C. tropicalis* is a human pathogenic ascomycete, closely related to the well-studied model fungal pathogen *C. albicans* (38). These two species diverged from their common ancestor ∼39 million years ago (39) and evolved with distinct karyotypes (21), having different phenotypic traits (40), and ecological niches (41). While *C. albicans* remains the primary cause of candidiasis worldwide, systemic ICU-acquired candidiasis is primarily (30.5-41.6%) caused by *C. tropicalis* in tropical countries including India (42), Pakistan (43), and Brazil (44). Moreover, the occurrence of drug resistance, particularly multidrug resistance, in *C. tropicalis* is on the rise (42, 45, 46). Therefore, relatively less-studied *C. tropicalis* is emerging as a major threat for nosocomial candidemia with 29-72% broad spectrum mortality rate (47). Fluconazole resistance in *C. albicans* can be gained due to segmental aneuploidy of Chr5 containing long IRs at the centromere, by the formation of isochromosomes (48), which was also identified in Chr4 with IRs at its centromere (49). All seven centromeres in *C. tropicalis* are associated with long IRs with the potential to form isochromosomes.

Since the mechanism of homology search during HR is positively influenced by spatial proximity and the extent of DNA sequence homology (3, 50), at least in the engineered model systems, it is expected that spatially clustered homologous DNA sequences undergo more translocation events than other loci. Although these factors were not shown to be involved in karyotypic rearrangements during speciation, a retrospective survey in light of spatial proximity and homology now offers a better explanation. For example, the bipolar to the tetrapolar transition of the mating type locus in the *Cryptococcus* species complex was associated with inter-centromeric recombination following pericentric inversion (51). Similar inter-centromeric recombination has been reported in the common ancestor of two fission yeast species, *Schizosaccharomyces cryophilus* and *Schizosaccharomyces octosporus* (16). These examples raise an intriguing notion that centromeres serve as sites of recombination, which may lead to centromere loss and/or the emergence of ENCs. This notion is supported by the fact that DSBs at centromeres following fusion of the acentric fragments to other chromosomes led to chromosome number reduction in *Ashbya* species (15) and *Malassezia* species (52). Genomic instability at the centromere can also lead to fluconazole resistance, as in the case of isochromosome formation on Chr5 of *C. albicans* (48). Additionally, breaks at the centromeres were reported to be associated with cancers in humans (53).

What would be the consequence of the spatial proximity of chromosomal regions with high DNA sequence homology in other domains of life? Inter-chromosomal contacts between chromosome pairs have been correlated with the number of translocation events in both naturally occurring populations and experimentally induced mammalian cells (54-63). It has been suggested that contacts between various chromosomal territories as well as their relative positions in the nucleus influence the sites and frequency of translocation events both in flies and mammals (58, 64-68). While centromeres remained clustered either throughout the cell cycle or most parts of it in many fungal species, such is not the case in metazoan cells. Nevertheless, one of the well-studied translocation events, Robertsonian translocation (RT) involving fusion between arms of two different chromosomes near a centromere, is the most frequently detected chromosomal abnormality in humans (69). The occurrence of RT was first reported in grasshoppers (70) and subsequently it has been implicated in the karyotype evolution in humans (69), mice (71, 72), and wheat (73). Moreover, RTs cause sterility in humans (74), often linked with the heterogeneity of carcinomas (75), and implicated in genetic disorders (76). Intriguingly, cytological and Hi-C based evidence (77) of spatial proximity (reviewed in (6)) among the repeat-associated centromere DNA sequences (78) in these species supports a possibility that RTs may have been guided by spatial proximity. Similarly, chromoplexy, involving a series of translocation events among multiple chromosomes without alterations in the copy number, was identified in prostate cancers (79, 80). Although fine mapping of translocation events at the repetitive regions in human cancer cells is challenging, the growing evidence that such events are associated with the formation of micronuclei (81) supports the idea that the spatial genome organization may influence chromoplexy as well (82).

The identification of HIR-associated putative centromeres in *C. parapsilosis, C. sojae*, and *C. viswanathii* supports the idea that the unique centromeres might have evolved from an ancestral HIR-associated centromere (83) (Figure 5A). While HIR-associated centromeres of *C. tropicalis, C. sojae*, and *C. viswanathii* form on different DNA sequences, a well-conserved IR-motif was identified in this study that is present in multiple copies on the centromeric IR sequences across these three species. Some centromeres in *C. albicans* carry chromosome-specific IRs but lack IR-motifs. Besides, *CaCEN5* IRs could not functionally complement the centromere function in *C. tropicalis* for the *de novo* CENP-A^Cse4^ recruitment. This indicates a possible role of the conserved IR-motifs on species-specific centromere function (21). Therefore, the loss of HIR-associated centromeres in *C. albicans* that are only epigenetically propagated (22) clearly shows how the ability of *de novo* establishment of kinetochore assembly in an ancestral lineage can be lost in a derived lineage. However, the mechanism through which IR-motifs may regulate centromere identity remains to be explored.

**Figure 5.**
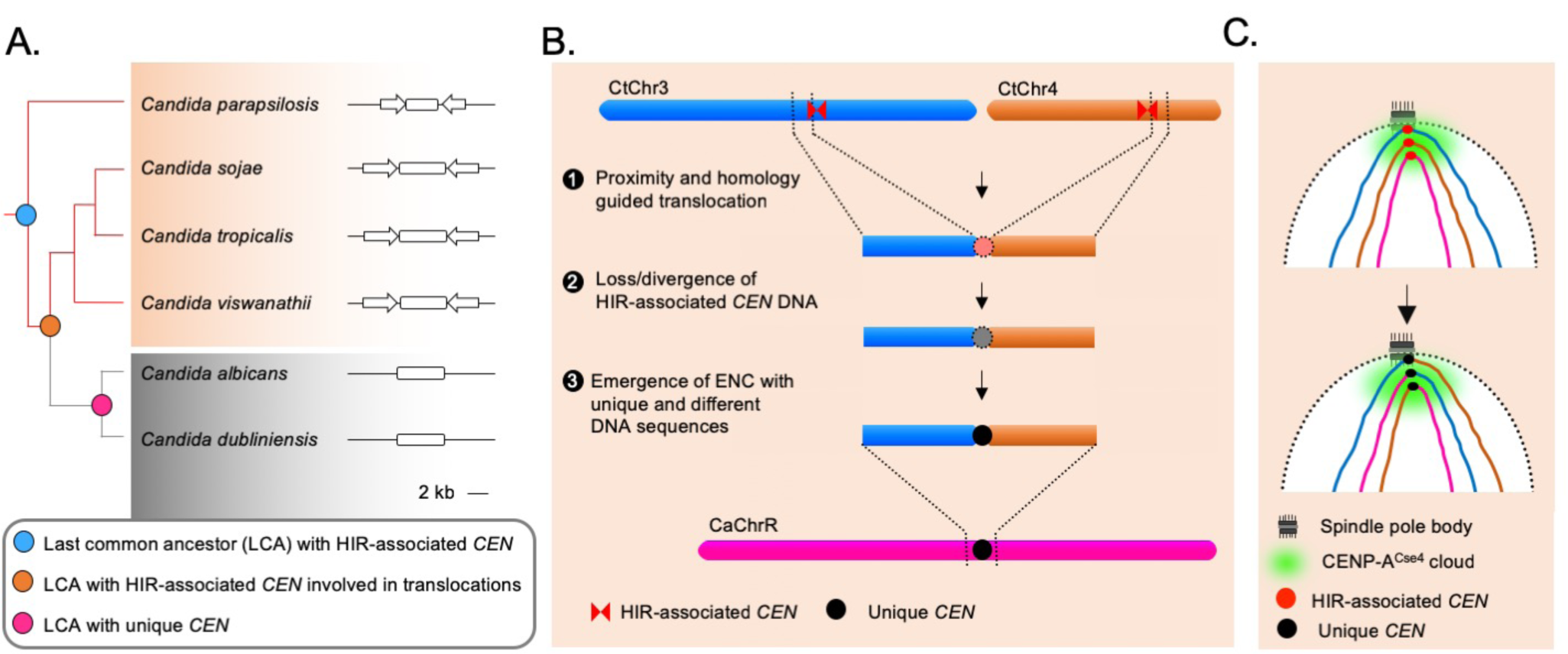
The spatial genome organization remained conserved in the CUG-Ser1 clade despite centromere type diversity. A. A maximum likelihood-based phylogenetic tree of closely related CUG-Ser1 species analyzed in this study. The centromere structure of each species is shown and drawn to scale. B. A model showing possible events during the loss of HIR-associated centromeres and emergence of the unique centromere type through inter-centromeric translocations possibly occurred in the common ancestor of *C. tropicalis* and *C. albicans.* The model is drawn to show translocation events involving two *C. tropicalis* chromosomes (CtChr3 and CtChr4) as representatives, which can be mapped proximal to the centromere on *C. albicans* ChrR (CaChrR) as shown in Figure 3F. C. Rabl-like chromosomal conformation is maintained despite inter-centromeric translocations that facilitated centromere type transition.

Loss of HIR-associated centromeres during inter-centromeric translocations or MIR must have been catastrophic for the cell, and the survivor was obligated to activate another centromere at an alternative locus. How is such a location determined? Artificial removal of a native centromere in *C. albicans* leads to the activation of a neocentromere (84, 85), which then becomes part of the centromere cluster (30). This evidence supports the existence of a spatial determinant, known as the CENP-A cloud or CENP-A-rich zone (84, 86), influencing the preferential formation of neocentromere at loci proximal to the native centromere (84, 87). We found that the unique and different centromeres of *C. albicans* are located proximal to the ORFs, which are also proximal to the centromeres in *C. tropicalis*. This observation indicates that the formation of the new centromeres in *C. albicans* may have been influenced by spatial proximity to the ancestral centromere cluster. However, new centromeres of *C. albicans* are formed on loci with completely unique and different DNA sequences. Similar to centromeres of *C. albicans*, centromere repositioning events may lead to the formation of ENCs, which are often associated with speciation in mammals (88, 89). It was found that the location of one centromere in horse varies across individuals (90, 91). Although, there are cases where ENCs formed without genomic rearrangements, the driving force facilitating centromere relocation was proposed to be associated with chromosomal inversion and translocation in certain cases (92). Because of these reasons, it may be logical to consider the centromeres of *C. albicans* as ENCs (Figure 5B). Intriguingly, even after the catastrophic chromosomal rearrangements, the ENCs in *C. albicans* remain clustered similar to *C. tropicalis* (Figure 5C). This observation identifies spatial clustering of centromeres as a matter of cardinal importance for the fungal genome organization.

## Methods

The strains, primers, and plasmids used in this study are listed in SI Appendix, Tables S8, S9, and S10, respectively. Details of all of the experimental procedures and data analysis are given in SI Methods. All sequencing data reported in the study and the genome assembly of *C. tropicalis* and *C. sojae* have been submitted to NCBI under the BioProject accession numbers PRJNA596050 and PRJNA604451.

## Acknowledgments

We thank all the members of KS laboratory and AS laboratory for stimulating discussions and critical reading of the manuscript. We acknowledge S. Sun and J. Heitman for helping with SMRT-seq of *C. tropicalis* at the PacBio sequencing facility at Duke University. We also thank A.I.S. Khalil for helping with CNAtra software for CNV analysis. Illumina sequencing experiments for the *C. sojae* genome were performed at Clevergene Biocorp, Bangalore, India. We also thank B. Suma for confocal microscopy, JNCASR. K.G. acknowledges Shyama Prasad Mukherjee Fellowship from Council of Scientific and Industrial Research (CSIR), Govt. of India [07/733(0181)/2013-EMR-I] and financial assistance from JNCASR. This project is supported by a grant (BT/PR27490/Med/29/1323/2018) from the Department of Biotechnology (DBT), Govt. of India to K.S. K.S. acknowledges TATA innovation fellowship (BT/HRD/35/01/03/2017) and Department of Biotechnology grant in Life Science Research, Education and Training at JNCASR (BT/INF/22/SP27679/2018). Intramural funding from JNCASR is acknowledged. This work is also supported by Nanyang Technological University’s Nanyang Assistant Professorship grant and Singapore Ministry of Education Academic Research Fund Tier 1 grant [RG39/18] to A.S.

## Author contributions

K.S. and A.S. supervised the study. K.S., A.S., K.G., and Y.C. conceived the idea and designed the experiments. K.G. performed N-gap filling, haplotype analysis, genome-wide synteny analyses, identification of the putative HIR-associated centromeres motif analysis, chromoblot analysis, Southern blotting, and subcellular localization experiments, Oxford Nanopore library preparation for *C. sojae* and generated its genome assembly. Y.C. analyzed 3C-seq data of *C. tropicalis* for chromatin contact map generation, aggregate signal analysis, and statistical comparison between telomeric interactions and bulk chromatin. With 3C-seq data YC validated joining of contigs and identified the CNVs in *the C. tropicalis* genome. B.C.T. and K.G. performed SMIS and Canu run, the scaffolding of telomeres, and Pilon polishing. R.M. and K.G. constructed aneuploid strains. S.R.B.M.M. performed the 3C-seq library preparation. C.O.B. and G.B. performed scaffolding of *C. tropicalis* genome in 16 contigs (Assembly B). K.G., K.S., Y.C., and A.S. wrote the manuscript and incorporated inputs from all of the authors. K.S. and A.S. edited the manuscript and provided the funding.

## References

1. Searle JB (1998) Speciation, chromosomes, and genomes. Genome Research 8(1):1–3.

2. Lee CS, et al.(2016) Chromosome position determines the success of double-strand break repair. Proceedings of the National Academy of Sciences of the United States of America 113(2):E146–154.

3. Agmon N, Liefshitz B, Zimmer C, Fabre E, & Kupiec M (2013) Effect of nuclear architecture on the efficiency of double-strand break repair. Nat Cell Biol 15(6):694–699.

4. Burgess SM & Kleckner N (1999) Collisions between yeast chromosomal loci in vivo are governed by three layers of organization. Genes Dev 13(14):1871–1883.

5. Piazza A, Wright WD, & Heyer WD (2017) Multi-invasions are recombination byproducts that induce chromosomal rearrangements. Cell 170(4):760–773 e715.

6. Muller H, Gil J, Jr., & Drinnenberg IA (2019) The impact of centromeres on spatial genome architecture. Trends Genet 35(8):565–578.

7. Clarke L & Carbon J (1980) Isolation of a yeast centromere and construction of functional small circular chromosomes. Nature 287(5782):504–509.

8. Mahtani MM & Willard HF (1990) Pulsed-field gel analysis of α-satellite DNA at the human X chromosome centromere: high-frequency polymorphisms and array size estimate. Genomics 7(4):607–613.

9. Navarro-Mendoza MI, et al.(2019) Early diverging fungus *Mucor circinelloides* lacks centromeric histone CENP-A and displays a mosaic of point and regional centromeres. Curr Biol 29(22):3791–3802 e3796.

10. Meraldi P, McAinsh AD, Rheinbay E, & Sorger PK (2006) Phylogenetic and structural analysis of centromeric DNA and kinetochore proteins. Genome Biol 7(3):R23.

11. Tromer EC, van Hooff JJE, Kops G, & Snel B (2019) Mosaic origin of the eukaryotic kinetochore. Proceedings of the National Academy of Sciences of the United States of America 116(26):12873–12882.

12. van Hooff JJ, Tromer E, van Wijk LM, Snel B, & Kops GJ (2017) Evolutionary dynamics of the kinetochore network in eukaryotes as revealed by comparative genomics. EMBO Reports 18(9):1559–1571.

13. Ekwall K (2007) Epigenetic control of centromere behavior. Annu Rev Genet 41(1):63–81.

14. Clarke L & Baum MP (1990) Functional analysis of a centromere from fission yeast: a role for centromere-specific repeated DNA sequences. Molecular and Cellular Biology 10(5):1863–1872.

15. Gordon JL, Byrne KP, & Wolfe KH (2011) Mechanisms of chromosome number evolution in yeast. PLoS Genet 7(7):e1002190.

16. Tong P, et al.(2019) Interspecies conservation of organisation and function between nonhomologous regional centromeres. Nature Communications 10(1):2343.

17. Kobayashi N, et al.(2015) Discovery of an unconventional centromere in budding yeast redefines evolution of point centromeres. Curr Biol 25(15):2026–2033.

18. Sanyal K, Baum M, & Carbon J (2004) Centromeric DNA sequences in the pathogenic yeast *Candida albicans* are all different and unique. Proceedings of the National Academy of Sciences of the United States of America 101(31):11374–11379.

19. Padmanabhan S, Thakur J, Siddharthan R, & Sanyal K (2008) Rapid evolution of Cse4p-rich centromeric DNA sequences in closely related pathogenic yeasts, *Candida albicans* and *Candida dubliniensis*. Proceedings of the National Academy of Sciences of the United States of America 105(50):19797–19802.

20. Kapoor S, Zhu L, Froyd C, Liu T, & Rusche LN (2015) Regional centromeres in the yeast *Candida lusitaniae* lack pericentromeric heterochromatin. Proceedings of the National Academy of Sciences of the United States of America 112(39):12139–12144.

21. Chatterjee G, et al.(2016) Repeat-associated fission yeast-like regional centromeres in the ascomycetous budding yeast *Candida tropicalis*. PLoS Genet 12(2):e1005839.

22. Baum M SK, Mishra PK, Thaler N, Carbon J. (2006) Formation of functional centromeric chromatin is specified epigenetically in *Candida albicans*. Proceedings of the National Academy of Sciences 103(40)(Oct 3):14877–14882.

23. Shen XX, et al.(2018) Tempo and mode of genome evolution in the budding yeast subphylum. Cell 175(6):1533–1545 e1520.

24. Butler G, et al.(2009) Evolution of pathogenicity and sexual reproduction in eight *Candida* genomes. Nature 459(7247):657–662.

25. Sexton T, et al.(2012) Three-dimensional folding and functional organization principles of the Drosophila genome. Cell 148(3):458–472.

26. Khalil AIS, Khyriem C, Chattopadhyay A, & Sanyal A (2020) Hierarchical discovery of large-scale and focal copy number alterations in low-coverage cancer genomes. BMC Bioinformatics 21(1):147.

27. Jones T, et al.(2004) The diploid genome sequence of *Candida albicans*. Proceedings of the National Academy of Sciences of the United States of America 101(19):7329–7334.

28. Duan Z, et al.(2010) A three-dimensional model of the yeast genome. Nature 465(7296):363–367.

29. Descorps-Declere S, et al.(2015) Genome-wide replication landscape of *Candida glabrata*. BMC Biol 13:69.

30. Burrack LS, et al.(2016) Neocentromeres provide chromosome segregation accuracy and centromere clustering to multiple loci along a *Candida albicans* chromosome. PLoS Genet 12(9):e1006317.

31. Sreekumar L, et al.(2019) Cis- and trans-chromosomal interactions define pericentric boundaries in the absence of conventional heterochromatin. Genetics 212(4):1121–1132.

32. Sreekumar L, et al.(2019) Orc4 spatiotemporally stabilizes centromeric chromatin. bioRxiv:465880, DOI: 465810.461101/465880.

33. Soderlund C, Bomhoff M, & Nelson WM (2011) SyMAP v3.4: a turnkey synteny system with application to plant genomes. Nucleic Acids Res 39(10):e68.

34. Grabherr MG RP, Meyer M, Mauceli E, Alföldi J, Di Palma F, Lindblad-Toh K. (2010) Genome-wide synteny through highly sensitive sequence alignment: Satsuma. Bioinformatics. 26(9):1145–1151.

35. Drillon G, Carbone A, & Fischer G (2014) SynChro: a fast and easy tool to reconstruct and visualize synteny blocks along eukaryotic chromosomes. PLoS One 9(3):e92621.

36. Ola M, et al.(2020) Polymorphic centromere locations in the pathogenic yeast Candida parapsilosis. bioRxiv:2020.2004.2009.034512.

37. Tsui CK, Daniel HM, Robert V, & Meyer W (2008) Re-examining the phylogeny of clinically relevant *Candida* species and allied genera based on multigene analyses. FEMS Yeast Res 8(4):651–659.

38. Legrand M, Jaitly P, Feri A, d’Enfert C, & Sanyal K (2019) *Candida albicans*: An emerging yeast model to study eukaryotic genome plasticity. Trends Genet 35(4):292–307.

39. Kumar S, Stecher G, Suleski M, & Hedges SB (2017) TimeTree: a resource for timelines, timetrees, and divergence times. Mol Biol Evol 34(7):1812–1819.

40. Cavalheiro M & Teixeira MC (2018) *Candida* Biofilms: threats, challenges, and promising strategies. Front Med (Lausanne) 5:28.

41. Pappas PG, Lionakis MS, Arendrup MC, Ostrosky-Zeichner L, & Kullberg BJ (2018) Invasive candidiasis. Nat Rev Dis Primers 4:18026.

42. Chakrabarti A, et al.(2015) Incidence, characteristics and outcome of ICU-acquired candidemia in India. Intensive Care Med 41(2):285–295.

43. Farooqi JQ, et al.(2013) Invasive candidiasis in Pakistan: clinical characteristics, species distribution and antifungal susceptibility. J Med Microbiol 62(Pt 2):259–268.

44. da Costa VG, Quesada RM, Abe AT, Furlaneto-Maia L, & Furlaneto MC (2014) Nosocomial bloodstream *Candida* infections in a tertiary-care hospital in South Brazil: a 4-year survey. Mycopathologia 178(3-4):243–250.

45. Xiao M, et al.(2015) Antifungal susceptibilities of *Candida glabrata* species complex, *Candida krusei, Candida parapsilosis* species complex and *Candida tropicalis* causing invasive candidiasis in China: 3 year national surveillance. J Antimicrob Chemother 70(3):802–810.

46. Goncalves SS, Souza ACR, Chowdhary A, Meis JF, & Colombo AL (2016) Epidemiology and molecular mechanisms of antifungal resistance in *Candida* and *Aspergillus*. Mycoses 59(4):198–219.

47. Lamoth F, Lockhart SR, Berkow EL, & Calandra T (2018) Changes in the epidemiological landscape of invasive candidiasis. J Antimicrob Chemother 73(suppl_1>):i4–i13.

48. Selmecki A, Forche A, & Berman J (2006) Aneuploidy and isochromosome formation in drug-resistant *Candida albicans*. Science 313(5785):367–370.

49. Todd RT, Wikoff TD, Forche A, & Selmecki A (2019) Genome plasticity in *Candida albicans* is driven by long repeat sequences. Elife 8:e45954.

50. Seeber A, Hauer MH, & Gasser SM (2018) Chromosome dynamics in response to DNA damage. Annu Rev Genet 52(1):295–319.

51. Wolfe K, et al.(2017) Fungal genome and mating system transitions facilitated by chromosomal translocations involving intercentromeric recombination. PLOS Biology 15(8):e2002527.

52. Sankaranarayanan SR, et al.(2020) Loss of centromere function drives karyotype evolution in closely related *Malassezia* species. eLife 9:e53944.

53. Barra V & Fachinetti D (2018) The dark side of centromeres: types, causes and consequences of structural abnormalities implicating centromeric DNA. Nat Commun 9(1):4340.

54. Arsuaga J, et al.(2004) Chromosome spatial clustering inferred from radiogenic aberrations. International journal of radiation biology 80(7):507–515.

55. Bickmore WA & Teague P (2002) Influences of chromosome size, gene density and nuclear position on the frequency of constitutional translocations in the human population. Chromosome research 10(8):707–715.

56. Branco MR & Pombo A (2006) Intermingling of chromosome territories in interphase suggests role in translocations and transcription-dependent associations. PLoS biology 4(5).

57. Canela A, et al.(2017) Genome organization drives chromosome fragility. Cell 170(3):507-521. e518.

58. Engreitz JM, Agarwala V, & Mirny LA (2012) Three-dimensional genome architecture influences partner selection for chromosomal translocations in human disease. PloS one 7(9).

59. Hlatky L, Sachs RK, Vazquez M, & Cornforth MN (2002) Radiation7induced chromosome aberrations: Insights gained from biophysical modeling. Bioessays 24(8):714–723.

60. Holley W, Mian I, Park S, Rydberg B, & Chatterjee A (2002) A model for interphase chromosomes and evaluation of radiation-induced aberrations. Radiation research 158(5):568–580.

61. Klein IA, et al.(2011) Translocation-capture sequencing reveals the extent and nature of chromosomal rearrangements in B lymphocytes. Cell 147(1):95–106.

62. Roukos V, Burman B, & Misteli T (2013) The cellular etiology of chromosome translocations. Current Opinion in Cell Biology 25(3):357–364.

63. Zhang Y, et al.(2012) Spatial organization of the mouse genome and its role in recurrent chromosomal translocations. Cell 148(5):908–921.

64. Aten JA, et al.(2004) Dynamics of DNA double-strand breaks revealed by clustering of damaged chromosome domains. Science 303(5654):92–95.

65. Foster HA, et al.(2013) Relative proximity of chromosome territories influences chromosome exchange partners in radiation-induced chromosome rearrangements in primary human bronchial epithelial cells. Mutation Research/Genetic Toxicology and Environmental Mutagenesis 756(1-2):66–77.

66. Savage JR (1998) A brief survey of aberration origin theories. Mutation Research/Fundamental and Molecular Mechanisms of Mutagenesis 404(1-2):139–147.

67. Savage JR (2000) Proximity matters. Science 290(5489):62–63.

68. Meaburn KJ (2016) Spatial genome organization and its emerging role as a potential diagnosis tool. Frontiers in genetics 7:134.

69. Therman E, Susman B, & Denniston C (1989) The nonrandom participation of human acrocentric chromosomes in Robertsonian translocations. Annals of human genetics 53(1):49–65.

70. Robertson WRB (1916) Chromosome studies. I. Taxonomic relationships shown in the chromosomes of *Tettigidae* and *Acrididae*: V7shaped chromosomes and their significance in *Acrididae, Locustidae*, and *Gryllidae*: chromosomes and variation. Journal of Morphology 27(2):179–331.

71. Castiglia R & Capanna E (2002) Chiasma repatterning across a chromosomal hybrid zone between chromosomal races of *Mus musculus domesticus*. Genetica 114(1):35–40.

72. Dumas D & Britton-Davidian J (2002) Chromosomal rearrangements and evolution of recombination: comparison of chiasma distribution patterns in standard and Robertsonian populations of the house mouse. Genetics 162(3):1355–1366.

73. Friebe B, Zhang P, Linc G, & Gill B (2005) Robertsonian translocations in wheat arise by centric misdivision of univalents at anaphase I and rejoining of broken centromeres during interkinesis of meiosis II. Cytogenetic and genome research 109(1-3):293–297.

74. Guichaoua M, et al.(1990) Infertility in human males with autosomal translocations: meiotic study of a 14; 22 Robertsonian translocation. Human genetics 86(2):162–166.

75. Hermsen M, et al.(2005) Centromeric chromosomal translocations show tissue-specific differences between squamous cell carcinomas and adenocarcinomas. Oncogene 24(9):1571–1579.

76. Mattei M, Souiah N, & Mattei J (1984) Chromosome 15 anomalies and the Prader-Willi syndrome: cytogenetic analysis. Human genetics 66(4):313–334.

77. Imakaev M, et al.(2012) Iterative correction of Hi-C data reveals hallmarks of chromosome organization. Nat Methods 9(10):999–1003.

78. Kalitsis P, Griffiths B, & Choo KA (2006) Mouse telocentric sequences reveal a high rate of homogenization and possible role in Robertsonian translocation. Proceedings of the National Academy of Sciences 103(23):8786–8791.

79. Zhang CZ, Leibowitz ML, & Pellman D (2013) Chromothripsis and beyond: rapid genome evolution from complex chromosomal rearrangements. Genes Dev 27(23):2513–2530.

80. Baca SC, et al.(2013) Punctuated evolution of prostate cancer genomes. Cell 153(3):666–677.

81. Crasta K, et al.(2012) DNA breaks and chromosome pulverization from errors in mitosis. Nature 482(7383):53–58.

82. Meaburn KJ, Misteli T, & Soutoglou E (2007) Spatial genome organization in the formation of chromosomal translocations. Seminars in cancer biology, (Elsevier), pp 80–90.

83. Coughlan AY, Hanson SJ, Byrne KP, & Wolfe KH (2016) Centromeres of the yeast *Komagataella phaffii (Pichia pastoris*) have a simple inverted-repeat structure. Genome Biology and Evolution 8(8):2482–2492.

84. Thakur J & Sanyal K (2013) Efficient neocentromere formation is suppressed by gene conversion to maintain centromere function at native physical chromosomal loci in *Candida albicans*. Genome Res 23(4):638–652.

85. Ketel C, et al.(2009) Neocentromeres form efficiently at multiple possible loci in *Candida albicans*. PLoS Genet 5(3):e1000400.

86. Fukagawa T & Earnshaw WC (2014) Neocentromeres. Curr Biol 24(19):R946–947.

87. Scott KC & Sullivan BA (2014) Neocentromeres: a place for everything and everything in its place. Trends Genet 30(2):66–74.

88. Rocchi M, Archidiacono N, Schempp W, Capozzi O, & Stanyon R (2012) Centromere repositioning in mammals. Heredity 108(1):59–67.

89. Stanyon R, et al.(2008) Primate chromosome evolution: ancestral karyotypes, marker order and neocentromeres. Chromosome Research 16(1):17–39.

90. Wade C, et al.(2009) Genome sequence, comparative analysis, and population genetics of the domestic horse. Science 326(5954):865–867.

91. Purgato S, et al.(2015) Centromere sliding on a mammalian chromosome. Chromosoma 124(2):277–287.

92. Schubert I (2018) What is behind “centromere repositioning”? Chromosoma 127(2):229–234.

93. Schindelin J, et al.(2012) Fiji: an open-source platform for biological-image analysis. Nature methods 9(7):676–682.

