## Supplemental Information for "Spatial inter-centromeric interactions facilitated the emergence of evolutionary new centromeres"

[1]Molecular Mycology Laboratory, Molecular Biology and Genetics Unit, Jawaharlal Nehru Centre for Advanced Scientific Research, Bangalore 560064, India; [2]School of Biological Sciences, Nanyang Technological University, 60 Nanyang Drive, Singapore 637551; [3]School of Biomolecular & Biomed Science, Conway Institute of Biomolecular and Biomedical Research, University College Dublin, Belfield, Dublin 4, Ireland. [4]Graduate School of Frontier Biosciences, Osaka University, Suita, Osaka 565 0871, Japan

*corresponding author

Kaustuv Sanyal

Amartya Sanyal

This PDF file includes:

Supplementary methods, references and figure legends

Figures S1 to S10

Tables S1 to S10

### Supplementary Methods

#### Media

*C. tropicalis* and *C. sojae* strains (Table S8) used in this study were grown in non-selective YPDU (2% dextrose, 2% peptone, 1% yeast extract, and 0.01% uracil), and incubated at 30°C at 180 rpm. For growing *C. albicans* strains, YPD media was supplemented with 0.1 mg/mL of uridine. The transformation of *C. tropicalis* was performed as described previously (1). The selection of transformants was based on prototrophy for the metabolic markers used. In the case of selection for the antibiotic marker (*CaSAT1*), conferring nourseothricin (NTC) resistance, growth media was supplemented with 100 µg/mL NTC (NTC; Werner Bioagents, CAS No. 96736-11-7). Recycling of the *CaSAT1* marker was done by growing the NTC resistant strains in YPMU (4% maltose, 2% peptone, 1% yeast extract, and 0.01% uracil) and segregants which are NTC sensitive were selected by patching them on YPDU and YPDU supplemented with NTC. For counter selection against *CaURA3*, the 5-Fluoroorotic Acid (5-FOA; Sigma-Aldrich, CAS No. 207291-81-4) was used at 1 mg/mL concentration.

#### Pulsed-field gel electrophoresis

*C. tropicalis* strain MYA-3404 and *C. albicans* strain SC5314 were grown until the exponential phase (~$2\times10^7$ cells/mL). Cells were washed with 50 mM EDTA and counted with a hemocytometer. Approximately $6\times10^8$ cells were used for the preparation of 1 mL genomic DNA plugs. The plugs were made according to the instruction manual protocol (Bio-Rad, Cat No. 170–3593) with CleanCut Agarose (0.6%) and the lyticase enzyme provided by the kit. A 0.6% pulsed field certified agarose gel was prepared using 0.5x TBE buffer (0.1 M Tris, 0.09 M Boric acid, 0.01 M EDTA, pH 8.0) and PFGE was performed on contour-clamped homogeneous electric field (CHEF) system using CHEF-DR II (Bio-Rad) module. The running conditions used were as follows: block-I at 100-200 s for 24 h at 4.5 V/cm/120°, block-II at 200-400 s for 48 h at 2.5 V/cm/120°, block-III at 600-800 s for 120 h at 2.5

V/cm/120°. The gel was stained with ethidium bromide (EtBr) and analyzed by Quantity One software (Bio-Rad).

**Indirect immunofluorescence microscopy**

Subcellular localization of Protein-A tagged CENP-$A^{Cse4}$ with DAPI (4′,6-diamidino-2-phenylindole) stained nuclear mass was performed in *C. tropicalis* strain CtKS102 following the method described previously for *C. albicans* (2). Asynchronously grown *C. tropicalis* cells were fixed with the 1/10$^{th}$ volume of formaldehyde (37%) for 1 h at room temperature. Antibodies used were diluted as follows: 1:1000 for rabbit anti-Protein A (Sigma, Cat No. P3775). The dilutions for secondary antibodies used were Alexa flour 568 goat anti-rabbit IgG (Invitrogen, Cat No. A11011) 1:1000. Antibody dilutions were prepared in 5% skimmed milk (HiMedia, Cat No. GRM1254) solution in 1x phosphate buffered saline (PBS) pH 7.4 (137 mM NaCl, 2.7 mM KCl, 10 mM $Na_2HPO_4$, 1.8 mM $KH_2PO_4$).

**Preparation of high molecular weight genomic DNA**

Briefly, 50 $OD_{600}$ equivalent (1 $OD_{600}$ = ~2×10$^7$ cells) cells were collected, washed with chilled 50 mM EDTA pH 8.0 and flash-frozen with liquid nitrogen. Next, the cell pellet was lyophilized. Then a volume equivalent to 5 mL of glass beads was added to the tube and vortexed till the pellet turns powdery. Then 20 mL Cetyltrimethyl ammonium bromide (CTAB) extraction buffer (100 mM Tris-HCl pH 7.5, 0.7 M NaCl, 10 mM EDTA, 1% CTAB powder, 1% 2-Mercaptoethanol) was added, and the tube was incubated at 65°C for ~30 min with occasional mixing by inverting the tube. Subsequently the tube was chilled on ice for 10 min, and the supernatant was transferred into another tube. An equal volume of chloroform was mixed with the supernatant gently inverting for 5 to 10 min. The mix was then centrifuged at 3200 rpm for 10 min, and the aqueous phase was carefully pipetted out using cut tips to a fresh tube. An equal volume of isopropanol was added into the supernatant and mixed gently until white thread-like structures appeared. The mix was incubated at −20°C for 1 h and centrifuged at 3200 rpm for 10 min to pellet the DNA. The pellet was washed twice with freshly prepared 70% ethanol and air-dried. The dried pellet was dissolved in 1 mL of 1x TE containing RNase A to a final concentration of 100 µg/mL and incubated at 37°C for 30 to 45 min. Sodium acetate solution was added into the mix to a final concentration of 0.5 M, and the solution was transferred to several 1.5 mL tubes in the aliquots of 0.4 mL each. An equal volume of isopropanol was added to each tube, mixed gently, and centrifuged at 13,000 rpm for 15 min. The supernatant was decanted, and the DNA pellet was washed with 70% ethanol. The pellet was air-dried and finally dissolved in 200 µL of 1x TE buffer. The quality of the isolated DNA was determined by performing PFGE analysis (switching time 1-25 s, at 5.8 V/cm/120° for 24 h, 1% agarose gel) on CHEF-DR II module (Bio-Rad).

**Oxford Nanopore sequencing of *C. sojae* strain NCYC-2607**

High molecular weight genomic DNA was isolated from yeast cells, and the average length of the DNA fragments of the genomic DNA was checked on a CHEF gel using a CHEF-DR II system (Bio-Rad). Next, the DNA sample was quantified by NanoDrop (ND-1000 Spectrophotometer, NanoDrop Technologies) and Qubit 3 fluorometer (Thermo Fisher Scientific) using dsDNA HS assay kit (Thermo Fisher Scientific, Cat No. Q33230). An appropriate amount of DNA was taken forward for library preparation as per the manufacturer's instructions using reagents included in SQK-LSK109 and EXP-NBD103/EXP-NBD104 kits. DNA samples were then pooled together on a single R9 flow-cell, and sequenced by the MinION system (Oxford Nanopore Technologies). The fragmentation step was skipped to retain the longer fragments. The raw reads were taken forward for base calling using Guppy version 3.1.5. A total of 92320 reads containing 530421800 bp were generated.

**Illumina sequencing of *C. sojae* strain NCYC-2607**

DNA was quantified by Qubit 3 fluorometer (Thermo Fisher Scientific) using a dsDNA HS assay kit (Thermo Fisher Scientific, Cat No. Q33230). Approximately 100 ng of intact DNA was enzymatically fragmented by targeting 250-500 bp fragment size. The DNA fragments with overhangs resulting from fragmentation were end-filled. The 3′ to 5′ exonuclease activity of end-repair mix removed the 3′ overhangs, and polymerase activity filled in the 5′ overhangs. To the blunt-ended fragments, adenylation was performed by adding a single 'A' nucleotide to the 3′ ends. To the adenylated fragments, loop adapters were ligated and cleaved with uracil-specific excision reagent enzyme. The sample was further purified using AMPure XP beads (Beckman Coulter, Cat No. A63880), and DNA was then enriched by PCR with six cycles using NEBNext Ultra II Q5 master mix (NEB, Cat No. M0544S), Illumina universal primers, and sample-specific indexed Illumina primers. The amplified products were cleaned up by using AMPure XP beads, and the final DNA library was eluted in 15 µL of 0.1x TE buffer. One µL of the library was used to quantify the DNA concentration by Qubit 3 fluorometer using the dsDNA HS reagent. The fragment analysis was performed on Agilent 2100 Bioanalyzer (Agilent, Model G2939B), by loading 1 µL of the library into Agilent DNA 7500 chip. In this experiment, we generated 3501768 paired-end reads of 2×301 bp length.

**_De novo_ genome assembly of _C. sojae_ strain NCYC-2607**

A total of 92320 reads containing 530421800 bp were used for the construction of a *de novo* assembly using Canu (3). Canu was run using default parameters in the trimming and the correction mode with '-genomeSize <15m>', which produced the genome assembly of *C. sojae* in 42 contigs. Next, to rectify the base-pair level errors, we performed five rounds of polishing of the contigs using Illumina reads with Pilon (4).

**SMRT sequencing on PacBio sequel system**

The genomic DNA fragments of ~20 kb length were size-selected and taken forward for library preparation using SMRTbell™ Template Prep Kit (Part No. 100-259-100). PacBio sequencing of the *C. tropicalis* MYA-3404 genome was performed by Sequel SMRT Cell 1M (Part No. 101-008-000) using Sequel™ Binding Kit 2.0 (Part no. 100-862-200) and SMRT Link version 5.0.1.9585. This run generated 996041 reads with an average read length of 5.8 kb.

**Construction of Assembly B**

Gepard (5) was used to generate dot matrix plots and identify areas of overlap between supercontigs. Supercontigs whose ends overlapped were identified and their sequences merged. Whole genome Illumina sequencing data of the *C. tropicalis* strain MYA-3404 were used to verify these predictions. We submitted the reads to NCBI under the BioProject accession number PRJNA604451.

**Construction of the _de novo_ SMRT assembly and contig stitching using SMIS**

The *de novo* SMRT assembly using 996041 PacBio raw reads was generated using Canu 1.6 (3). The program was run in the trimming and correction mode with the ‘-pacbio-raw <input.fastq>’ option that produced 135 contigs. For stitching the contigs from Assembly B using the PacBio raw reads, we used Single Molecular Integrative Scaffolding (https://github.com/fg6/smis) with the default options, to get a 12-contig assembly (Assembly C). Details of the assemblies produced by Canu and SMIS are presented in Table S3**.**

**Filling N-gaps**

The *de novo* SMRT contigs were used to fill the existing N-gaps in Assembly A. We used 500 bases upstream, and downstream regions of the N-gaps as queries against a custom BLAST (6) database generated using Geneious® software from the *de novo* assembled contigs and filled these N-gaps upon the mapping of upstream and downstream query sequences on the same contig with 100% coverage and more than 95% identity. Using this approach, we filled 78 out of 104 gaps leaving 26 gaps on seven chromosomes (Table S2, Figure S3A). We suspected that the remaining gaps were repetitive regions in the genome as immediate flanking regions identified multiple hits. To avoid this, we used a second strategy in which we used a 1-kb query sequence from either 10 kb upstream or downstream region of the N-gap, and performed a BLAST analysis against the *de novo* contigs generated using FALCON (7). All the remaining 26 gaps could be filled using this strategy (Table S2, Figure S3B). Further, to validate our claim, we confirmed the mapping of the Illumina and PacBio reads over the newly filled sequence.

**Assembly of sub-telomeric regions**

To assemble the sub-telomere regions, we performed a BLAST search using the terminal 5000 bp sequence of each chromosome as queries against the *de novo* SMRT contigs and identified the contigs containing the 23-bp telomeric repeats specific for *C. tropicalis* (5′-TGATCGTGACATCCTTACACCAA-3′) as reported previously (8). Schematic of the sub-telomere scaffolding has been shown (Figure S3C).

**Mapping of the orphan haplotigs using the *de novo* SMRT assembly**

Canu is a diploid-aware genome assembler (3), which generates two contigs from a heterozygous locus. Therefore, we used the Canu generated contigs (SMRT assembly) to map the orphan haplotigs as heterozygous regions of the genome (see Figure S1H). Heterozygosity of the orphan haplotigs was demonstrated by the Illumina read coverage (Figure S2B). For this analysis, the 3C-seq reads were mapped on the OHs and a control locus of Chr1 using Bowtie2 (9). The number of mapped reads were counted using the bamCoverage utility from deepTools2 (10) and plotted using boxplotR (11).

**Pilon polishing of the genome assembly**

The final telomere-to-telomere assembled chromosomes were polished through Pilon (4) using the Illumina reads obtained from the 3C-seq experiment. Pilon corrected base-pair level assembly errors and validated 99.5-99.8% bases of the seven chromosomes. The polishing step was repeated six times when the improvement stalled.

**Construction of aneuploids for confirmation of heterozygosity of the OHs**

We constructed *C. tropicalis* strains monosomic for Chr5 and used them to demonstrate that loss of one homolog of Chr5 leads to loss of one of the two alleles of the orphan contigs: contig14 and contig16, that are mapped on Chr5. Since the *sch9* mutants in *C. albicans* were viable but lost chromosomes at a significantly higher rate than the wild-type (12), we adopted the same strategy to delete both copies of *SCH9* homologs in *C. tropicalis.* Next, a reporter strain was created in this *sch9* mutant strain background of *C. tropicalis* to assay for loss of a Chr5 homolog. These strains (2n-1) that lacked one homolog of Chr5 were used to confirm the presence of heterozygosity of orphan haplotigs (OHs) of CtChr5.

a. **Deletion of *SCH9* in *C. tropicalis***

The *SCH9* homolog in *C. tropicalis* was identified in a BLAST search using *CaSCH9* as the query sequence against the *C. tropicalis* proteome. A putative homolog of *SCH9* was located on Chr1:1994521-1996662 and encoded by the Crick strand. A deletion cassette (pKG1) for double homologous recombination-mediated deletion of *SCH9* ORF was constructed by cloning upstream and downstream homology regions in pSFS2a plasmid (13). This construct was transformed into CtKS102 for the deletion of both copies of *SCH9* ORF by recycling the *CaSAT1* marker after the deletion of the first copy of *SCH9* gene. Independent colonies of the *sch9*/*sch9* null mutant strain (CtKG001) were confirmed using Southern hybridization (Figure S2C-D). Primers used in this study are mentioned in Table S9.

**b. Construction of a reporter strain for construction of strains with Chr5 monosomy by integration of *URA3* on Chr5**

Upstream and downstream homology regions of the target intergenic locus (Chr5_497_kb) in Chr5 were amplified, and cloned into pBSCaURA3 plasmid (1) to construct pKG2 (Table S10). This cassette was released by restriction digestion with BamHI and ApaI and transformed into the *sch9* mutant strain CtKG001 to construct the reporter strain CtKG002. Similarly, we integrated *CaURA3* into the target intergenic locus (Chr5_497_kb) of CtKS102 to create a control strain CtKG003. In both the strains (CtKG002 and CtKG003) the short arm (5′ end) of one of the two homologs is marked with *CaURA3* marker and the long arm (3′ end) carries the heterozygous *MTL* locus (*MTL***a** or *MTLα*) with two distinct alleles present on two homologs. Concomitant loss of one of the *MTL* alleles together with *CaURA3* marker would indicate loss of one homolog of Chr5.

**c. Isolation and confirmation of the 2n-1 aneuploids for Chr5**

Different cell numbers ($10^5$, $10^4$, $10^3$, and $10^2$) of the reporter strain (CtKG002) and the wild-type control strain (CtKG003) were plated on complete media (CM) + 5-FOA and incubated for 48-72 h at 30°C. Multiple $FOA^R$ colonies appeared for CtKG003 strain but no colonies appeared for the control strain (CtKG003). The colonies were then patched on YPDU and CM-URA plates for confirmation of the loss of the *CaURA3* marker. Next, PCR was performed to confirm the loss of one of the *MTL* loci (*MTL***a** or *MTLα)* in these colonies using a multiplex PCR strategy described previously (Figure S2G) (14).

**Library preparation and sequencing of the library DNA for chromosome conformation capture (3C-seq)**

Wild-type *C. tropicalis* strain MYA-3404 was cultured in non-selective YPDU media and 500 $OD_{600}$ equivalent cells were harvested for crosslinking. The cells were cross-linked with formaldehyde to a final concentration of 1.5% for 10 min and the cross-linking reaction was quenched by adding glycine to a final concentration of

400 mM. The crosslinked cells were centrifuged and the cell pellet was stored at -80°C till further use.

For making the 3C library of *C. tropicalis*, the cross-linked cell pellet was first resuspended in 5 mL of ice-cold 1x NEBuffer™ DpnII (50 mM Bis-Tris-HCl, 100 mM NaCl, 10 mM MgCl2, 1 mM DTT; pH 6 @ 25°C) and then lysed by liquid nitrogen grinding in a chilled mortar using a pestle to a fine powder. The powdered sample was scraped using a spatula into a pre-chilled tube and resuspended in 15 mL cold 1x NEBuffer™ DpnII. Cell lysate containing ~3×$10^8$ cells (4 mL) was processed for 3C library preparation. This lysate was centrifuged and the pellet was resuspended in 1.5 mL of cold 1x NEBuffer™ DpnII and then aliquoted equally into four 1.5 mL microcentrifuge tubes. Next, the chromatin was solubilized by adding SDS to a final concentration of 0.1% in each microcentrifuge tube and the sample was incubated at 65°C for exactly 10 min. The reaction was quenched by adding 45 µL of 10% Triton X-100 per tube with gentle mixing. Chromatin was then digested with 750 units of DpnII (NEB, Cat No. R0543M; 50,000 units/mL) per tube and incubated at 37°C overnight with gentle agitation (300 rpm) on a heating block. Next day, the restriction enzyme was heat-inactivated at 65°C for 20 min. The digested chromatin fraction in each tube was ligated with 50 U of T4 DNA ligase (Invitrogen Cat No.15224090; 1 U/µL) at 16°C for 6 h in a diluted condition (reaction volume 8 mL) to favor intra-molecular ligation of cross-linked restriction fragments. Reverse cross-linking was performed by adding 100 µL of 10 mg/mL Proteinase K (Invitrogen Cat No.25530031) per tube and incubating at 65°C overnight. Next, DNA, which constitutes the 3C library, was purified using conventional phenol-chloroform extraction and concentrated using Amicon Ultra-0.5 mL 30K centrifugal filters. About 1 µg of 3C library was used for size selection using Agencourt AMPure XP beads (Beckman Coulter) to select DNA fragments of 500-700 bp in length. The paired-end NGS library was prepared using NEBNext Ultra II kit, and sequencing was carried out using the Illumina HiSeq 2500 2×101 bp platform by a third party service provider.

**3C-seq data analysis**

FASTQ files containing ~75 million 2×101 bp paired-end 3C-seq reads were initially processed using hiclib package (http://mirnylab.bitbucket.org/hiclib/) (17). The resultant genome-wide chromatin interaction matrix was converted to a contact probability matrix. Codes associated with the downstream analysis could be found at the Github repository (https://github.com/Yao-Chen/candida-tropicalis-analysis).

**a. Mapping of reads and generation of the contact probability matrix**

First, two sides of paired-end reads were separated and iteratively aligned to Assembly2020 using Bowtie2 (9, 15) with the '--very-sensitive' option. The iteration started from the first 20 bases of each read and continued with an increment of 5 bases in the subsequent iteration. Next, the aligned read pairs whose both sides had MAPQ score ≥1 were processed through the fragment filter, where self-circles, dangling ends, extra-dangling ends (maximum molecular length = 500), error pairs and PCR duplicates were excluded from downstream analysis. A genome-wide interaction matrix was generated using the remaining unique valid pairs (bin size = 2-10 kb). The bin filter removed bins with <50% sequence information in the reference genome and 1% bins with low read coverage. The matrix was iteratively corrected for biases and eventually converted to a contact probability matrix where the sum of each row/column approximates 1.

**b. Aggregate signal analysis**

Sub-matrices for all combinations of centromere-centromere interactions across different chromosomes were extracted from the genome-wide contact probability matrix. Genomic loci containing mid-points of centromeres were first aligned and then all the sub-matrices were stacked on top of each other and averaged. Similar analysis was performed for all telomere-telomere interactions (both intra and interchromosomal telomeric interactions) where sub-matrices for telomere-telomere interactions across all chromosomes were extracted, stacked and averaged.

**c. Analysis of telomeric interactions**

To investigate interchromosomal telomeric interactions, a histogram of all interchromosomal interactions (excluding zero values) were generated (bulk chromatin). Mean contact probabilities of all interchromosomal interactions as well as all interchromosomal telomeric interactions were computed, where 5′ and 3′ end bins on each chromosome were taken as telomeric bins. Similarly, a histogram of all intrachromosomal long-range interactions (excluding zero values) was generated (bulk chromatin), where long-range interactions were defined as interactions between loci separated at a distance of >100 kb. Intrachromosomal telomeric interactions were taken as interactions between two loci that were close to two telomeres of an individual chromosome respectively (sum of distances between each locus to the nearest telomere is ≤10 kb). Mann-Whitney U test was used to compare interchromosomal or intrachromosomal telomeric interactions to the bulk chromatin.

**d. Contig scaffolding**

3C-seq reads were aligned to contig sequences and contact probability matrix was generated as described above. The 3C profile of a bin was plotted using values in a single row from the contact probability matrix. It is well-established that contact frequency generally shows a distance-dependent decay (16). Therefore, the connectivity between two contigs can be inferred by investigating the contact probabilities between the terminal bin of a contig and loci on the other contig.

**Identification of SNPs, indels and CNVs**

**a. Detection of SNPs and indels**

The SNPs and indels were identified using GATK software (17) with the paired-end Illumina reads obtained from the 3C-seq experiment in a 12 cores Ubuntu 16.4 system with 96 GB memory. Briefly, the 3C-seq reads were mapped to Assembly2020 using Bwa-mem (18) paired-end alignment mode following sorting of the resulting SAM file with Picard (https://broadinstitute.github.io/picard/), SAM to BAM conversion using SAMtools (19), and duplicate marking using ‘MarkDuplicates’ utility of Picard. Next, we used GenomeAnalysisTK.jar (version 3.8.0) to call the variants with ‘-ploidy 2’ option, SNPs were extracted, filtered with ‘--filterExpression 'QD < 2.0 || FS > 60.0 || MQ < 40.0 || MQRankSum < -12.5 || ReadPosRankSum < -8.0 || SOR > 4.0' --filterName "basic_snp_filter"’ option following base quality score recalibration. Similarly indels were extracted, filtered with ‘--filterExpression 'QD < 2.0 || FS > 200.0 || ReadPosRankSum < -20.0 || SOR > 10.0' --filterName "basic_indel_filter"’ option following base quality score recalibration. The data tracks were visualized using IGV (20) and presented using Circa software.

**b. Read coverage plot and CNV detection**

To generate a genome-wide read coverage plot, the 3C-seq reads were mapped to Assembly2020 using Bowtie2 (9) paired-end alignment mode with ‘--end-to-end’ and ‘--very-sensitive’ option. The resultant SAM file was converted to BAM format and sorted using SAMtools (19). Next, the mapped reads were counted using deepTools2 (10) bamCoverage utility with the BPM normalization method, and the resulting BED file was used for downstream calculations or visualization in IGV (20). To detect CNVs throughout the *C. tropicalis* genome, the sorted BAM file, generated from coverage analysis above, was processed by ‘CNAtraLite’ option of CNAtra tool (https://github.com/AISKhalil/CNAtra) (21). where the MAPQ filter was disabled. The read depth signal (bin size = 1 kb) and the estimated copy numbers were then plotted for each chromosome by ‘CNVsTrackPlot’ function of ‘CNAtraLite’. In this analysis, regions whose estimated copy numbers are <1.5 or >2.5 are considered as CNVs.

**Haplotype analysis**

The FALCON, FALCON-Unzip (7), and FALCON-Phase (22) from the pb-assembly suite were run locally in a 12 core Ubuntu 16.4 system with 96 GB memory according to the instruction provided (https://github.com/PacificBiosciences/pb-assembly). The configuration files used for running FALCON, FALCON-Unzip and FALCON-Phase will be available upon request. Briefly, FALCON was run using modified fcrun.cfg with the input option ‘pa_DBdust_option=true’, and ‘pa_fasta_filter_option=streamed-internal-median’. Next, data partitioning was performed with ‘pa_DBsplit_option=-x500 -s100’ and ‘ovlp_DBsplit_option=-x500 -s100’, repeat masking was performed using ‘pa_HPCTANmask_option = -k18 -h480 -w8 -e.8 -s100’, ‘pa_HPCREPmask_option = -k18 -h480 -w8 -e.8 -s100’, and ‘pa_REPmask_code=0,300;0,300;0,300’ options. Preassembly was generated using the following parameters: ‘genome_size=15000000’, ‘seed_coverage=20’, ‘length_cutoff=100’, ‘pa_HPCdaligner_option=-v -B128 -M24’, ‘pa_daligner_option=-e.8 -l1000 -k18 -h480 -w8 -s100 -T10’, ‘falcon_sense_option=--output-multi --min-idt 0.70 --min-cov 2 --max-n-read 1800’, ‘falcon_sense_greedy=False’. Next, Pread overlapping was performed using ‘ovlp_daligner_option=-e.96 -l1000 -k24 -h1024 -w6 -s100’, and ‘ovlp_HPCdaligner_option=-v -B128 -M24’. Next, the final assembly was generated using ‘overlap_filtering_setting=--max-diff 100 --max-cov 100 --min-cov 2’ and ‘length_cutoff_pr=500’. Next, phasing of haplotypes was performed using FALCON-Unzip and FALCON-Phase as described (https://github.com/PacificBiosciences/pb-assembly).

**Assessment of the genome assembly completeness using BUSCO**

BUSCO (23) version 3.0.2 was run against ascomycota_odb9 database using the following script: python ./scripts/run_BUSCO.py -i genome.fasta -o BUSCO_output -l /Path_to_llineage_dir/ -m genome -c 1 -sp candida_tropicalis.

**Synteny analysis**

Genome-wide synteny analysis was performed using Symap (24) with the parameters as (a) Min. dots 3 (minimum number of anchors required to define a synteny block), (b) top N 2 (retain the top N hits for every sequence region, as well as all hits with score at least 80% of the Nth), (c) BLAT args: '-minScore=30 -minIdentity=70 -tileSize=10 -qMask=lower -maxIntron=10000'. The Satsuma synteny and Synchro software were run using default parameters. For the custom approach to map the interchromosomal synteny breakpoints (ICSBs), first, the single-copy orthologs were identified using OrthoFinder (25), then the corresponding genomic coordinates of the ortholog pairs were sorted and the ICSBs were identified. For comparing the FALCON generated contigs with the Assembly2020 chromosomes, the dot-plot between the two assemblies was generated using the default options of Symap version 4.2.

**Identification of the putative centromeres in the members of CUG-Ser1 clade**

The putative centromeres of *C. sojae* and *C. viswanathii* were identified as HIR-associated intergenic regions syntenic to centromeres of *C. tropicalis* centromeres. Briefly, the genomic loci in *C. sojae* and *C. viswanathii,* which are syntenic to the centromeres of *C. tropicalis* were scanned for the presence of inverted repeats falling in ORF-free regions using YASS (26) with the default parameters. Pair-wise alignments between seven random genomic loci of ~10 kb length, LR, CC, or RR DNA elements were performed using Clustal Omega (27). Synteny dot-plot analysis for centromere DNA sequences including the flanking ORF-free region in *C. albicans, C. dublininensis* and the HIR sequences of *C. tropicalis, C. sojae, C. viswanathii,* and *C. parapsilosis* was generated using Gepard (5) by running it in the simple mode with default parameters. The IR sequences from centromeres of *C. tropicalis* and the putative centromeres of *C. sojae* and *C. viswanathii* were analysed to identify the presence of conserved motifs using motif discovery tool MEME following the default parameters with 'ZOOPS: zero or one site per sequence' as the motif site distribution algorithm, and maximum motif width set to 12 bp. Next, we scanned for the presence of IR-motifs across the chromosomes including centromere DNA and flanking ORF-free regions in *C. albicans, C. dubliniensis,* and putative centromeres of *C. parapsilosis* using FIMO with default parameters (28).

**Construction of the phylogenetic tree**

The publicly available genomes and the protein fasta files (when available) of *C. albicans* (ASM254v2), *C. dubliniensis* (ASM2694v1), *C. viswanathii* (ASM332773v1), and *C. parapsilosis* (ASM18276v2) were downloaded from NCBI database. The protein fasta files for *C. tropicalis* and *C. sojae* were generated using Augustus *ab initio* protein prediction software and the python script getAnnoFasta.py (29). Because of the partially diploid nature of *C. viswanathii* genome assembly, the duplicated contigs, that carried >100 kb of DNA sequence on another contig, were identified from dot-plot analysis (self) using Symap (24), and excluded from analysis. The protein fasta files were then used as input files for running OrthoFinder V2.3.1 (25). OrthoFinder was run using the default parameters except the -M msa option for the construction of maximum-likelihood trees using MAFFT (30) and FastTree (31). The tree topology was visualized using Evolview (32).

**References:**


1. Chatterjee G*, et al.* (2016) Repeat-associated fission yeast-like regional centromeres in the ascomycetous budding yeast *Candida tropicalis*. *PLoS Genet* 12(2):e1005839.
2. Sanyal K & Carbon J (2002) The CENP-A homolog CaCse4p in the pathogenic yeast *Candida albicans* is a centromere protein essential for chromosome transmission. *Proceedings of the National Academy of Sciences of the United States of America* 99(20):12969-12974.
3. Koren S*, et al.* (2017) Canu: scalable and accurate long-read assembly via adaptive k-mer weighting and repeat separation. *Genome Res* 27(5):722-736.
4. Walker BJ*, et al.* (2014) Pilon: an integrated tool for comprehensive microbial variant detection and genome assembly improvement. *PLoS One* 9(11):e112963.
5. Krumsiek J, Arnold R, & Rattei T (2007) Gepard: a rapid and sensitive tool for creating dotplots on genome scale. *Bioinformatics* 23(8):1026-1028.
6. Altschul SF, Gish W, Miller W, Myers EW, & Lipman DJ (1990) Basic local alignment search tool. *J Mol Biol* 215(3):403-410.
7. Chin CS*, et al.* (2016) Phased diploid genome assembly with single-molecule real-time sequencing. *Nat Methods* 13(12):1050-1054.
8. Butler G*, et al.* (2009) Evolution of pathogenicity and sexual reproduction in eight *Candida* genomes. *Nature* 459(7247):657-662.

9. Langmead B & Salzberg SL (2012) Fast gapped-read alignment with Bowtie 2. *Nat Methods* 9(4):357-359.
10. Ramirez F*, et al.* (2016) deepTools2: a next generation web server for deep-sequencing data analysis. *Nucleic Acids Res* 44(W1):W160-165.
11. Spitzer M, Wildenhain J, Rappsilber J, & Tyers M (2014) BoxPlotR: a web tool for generation of box plots. *Nat Methods* 11(2):121-122.
12. Varshney N*, et al.* (2015) A surprising role for the Sch9 protein kinase in chromosome segregation in *Candida albicans*. *Genetics* 199(3):671-674.
13. Reuss O, Vik A, Kolter R, & Morschhauser J (2004) The *SAT1* flipper, an optimized tool for gene disruption in *Candida albicans*. *Gene* 341:119-127.
14. Porman AM, Alby K, Hirakawa MP, & Bennett RJ (2011) Discovery of a phenotypic switch regulating sexual mating in the opportunistic fungal pathogen *Candida tropicalis*. *Proceedings of the National Academy of Sciences of the United States of America* 108(52):21158-21163.
15. Baca SC*, et al.* (2013) Punctuated evolution of prostate cancer genomes. *Cell* 153(3):666-677.
16. Dekker J, Rippe K, Dekker M, & Kleckner N (2002) Capturing chromosome conformation. *Science* 295(5558):1306-1311.
17. McKenna A*, et al.* (2010) The Genome Analysis Toolkit: a MapReduce framework for analyzing next-generation DNA sequencing data. *Genome Res* 20(9):1297-1303.
18. Li H (2013) Aligning sequence reads, clone sequences and assembly contigs with BWA-MEM. *arXiv preprint arXiv:1303.3997*.
19. Li H*, et al.* (2009) The sequence alignment/map format and SAMtools. *Bioinformatics* 25(16):2078-2079.
20. Robinson JT*, et al.* (2011) Integrative genomics viewer. *Nat Biotechnol* 29(1):24-26.
21. Khalil AIS, Khyriem C, Chattopadhyay A, & Sanyal A (2020) Hierarchical discovery of large-scale and focal copy number alterations in low-coverage cancer genomes. *BMC Bioinformatics* 21(1):147.
22. Kronenberg ZN*, et al.* (2018) FALCON-Phase: Integrating PacBio and Hi-C data for phased diploid genomes. *bioRxiv*:327064.
23. Simao FA, Waterhouse RM, Ioannidis P, Kriventseva EV, & Zdobnov EM (2015) BUSCO: assessing genome assembly and annotation completeness with single-copy orthologs. *Bioinformatics* 31(19):3210-3212.
24. Soderlund C, Bomhoff M, & Nelson WM (2011) SyMAP v3.4: a turnkey synteny system with application to plant genomes. *Nucleic Acids Res* 39(10):e68.
25. Emms DM & Kelly S (2015) OrthoFinder: solving fundamental biases in whole genome comparisons dramatically improves orthogroup inference accuracy. *Genome Biol* 16:157.
26. Noe L & Kucherov G (2005) YASS: enhancing the sensitivity of DNA similarity search. *Nucleic Acids Res* 33(Web Server issue):W540-543.
27. Sievers F & Higgins DG (2014) Clustal Omega, accurate alignment of very large numbers of sequences. *Multiple sequence alignment methods*, (Springer), pp 105-116.
28. Bailey TL*, et al.* (2009) MEME SUITE: tools for motif discovery and searching. *Nucleic Acids Res* 37(Web Server issue):W202-208.

29. Stanke M MB (2005) AUGUSTUS: a web server for gene prediction in eukaryotes that allows user-defined constraints. *Nucleic Acids Research* 33(suppl_2)(Jul 1):W465-467.
30. Katoh K, Misawa K, Kuma K, & Miyata T (2002) MAFFT: a novel method for rapid multiple sequence alignment based on fast Fourier transform. *Nucleic Acids Res* 30(14):3059-3066.
31. Price MN, Dehal PS, & Arkin AP (2010) FastTree 2-approximately maximum-likelihood trees for large alignments. *PLoS One* 5(3):e9490.
32. Subramanian B, Gao S, Lercher MJ, Hu S, & Chen W-H (2019) Evolview v3: a webserver for visualization, annotation, and management of phylogenetic trees. *Nucleic acids research* 47(W1):W270-W275.

### Supplementary figure legends:

**Figure S1. Schematic of the strategies used for construction of the gapless chromosome-level assembly of *C. tropicalis*.**
A - D. The outline of the steps followed for the construction of the genome assembly. A. Major steps followed for 3C-sequencing in this study were (I) crosslinking, (II) restriction digestion, and (III) ligation, library preparation, and sequencing. B. A cartoon explaining the use of contact probability values for establishing contiguity between two DNA fragments. Pink arrows denote coordinates of the anchor bin, with respect to which the contact probability scores (represented as the red dots) are determined. C. Long reads generated by SMRT-seq were used to construct *de novo* genome assemblies using Canu as well as the FALCON pipeline. D. Use of *de novo* contigs to fill N-gaps, finding the alleles of the orphan contigs in the genome and scaffolding of the sub-telomeres. E. The 3C profile (bin size = 10 kb) of 3′-terminal bin of contig6 (anchor; gray vertical line) showing its contact probabilities (blue dots) with bins on contig5 and contig6. F. The 3C profile (bin size = 10 kb) of 3′ terminal bin of contig5 (anchor; gray vertical line) showing its contact probabilities (blue dots) with bins on contig5 and contig6. G. A schematic representation of chromosome 2 assembled by fusion of contig5 and contig6 in a tail-to-tail orientation based on the 3C profile results. H. Orphan contigs OHs are mapped to the chromosomes by BLAST analysis of the ORFs, which are located on the Canu assembled *de novo* contigs. The allelic difference in the OH loci is depicted by color-coded ORFs (orange and green arrows).

**Figure S2. Orphan contigs are alleles in the diploid genome of *C. tropicalis.***

A. IGV track images showing the coverage of 3C-seq data on the *y*-axis (number of reads mapped per bin for each million of the total reads) over the orphan contigs and a control locus from Chr1. B. Violin plots showing the distribution of read coverage of 3C-seq data across the orphan contigs (bin size = 5 bp) and the control region on Chr1 generated using deepTools2 bamCoverage script. C. Schematic showing the positions of HindIII sites (vertical black lines) and the length of the expected bands detectable by the probe used (red bars) for Southern hybridization to confirm *sch9* deletion strains (CtKG001). D. Phosphorimage of the blot for confirmation of *sch9Δ/sch9Δ* mutant strains. E. Schematic showing various number of cells ($10^5$, $10^4$, $10^3$, and $10^2$) of CtKG002 (left) and CtKG003 (right) plated on CM+FOA plate. The plate image showing appearance of FOA$^R$ colonies in CtKG002 but not in CtKG003 strain. F. The FOA$^R$ colonies thus obtained were picked up, patched on CM-URA (left), and YPDU media (right) and imaged after 48 h of growth at 30°C . G. The EtBr stained gel image for multiplex PCR products to detect the loss of *MTL***a** or *MTLα* alleles in the FOA$^R$ colonies along with the wild-type control (Primers are listed on Table S9). H. and I. Experimental validation of the allelic nature of contig16 and contig14, respectively, using Southern blot analysis. The length of restriction fragments polymorphisms between the alleles after digesting with ClaI and EcoRI (restriction enzyme sites are indicated using blue arrowheads) are graphically represented for contig16 and contig14, respectively. The lanes in gels represent the wild-type MYA-3404 (2$^{nd}$ lane) and the monosomic aneuploid strains (CtKG101 - 105) where one homolog of Chr5 is absent. The probes used in this experiment are denoted using red bars.

**Figure S3. Schematic outline of the strategy followed for N-gap filling and scaffolding of sub-telomeres.**

A - B. Strategy-I and strategy-II (Methods) for filling N-gaps without flanking repeats or with flanking repeats, respectively. Repeats are presented as black arrows. C. Schematic for scaffolding of sub-telomeres using the *de novo* assembled contigs.

**Figure S4. Identification of CNVs in the *C. tropicalis* strain MYA-3404**

A. EtBr stained gel images and phosphorimages obtained from Southern hybridization experiments using a centromere-proximal (Probe A) and a centromere-distal probe (Probe B) from Chr4 (Table S9). B. Violin plots for the number of reads mapped per bin (2 bp) on ChrR (excluding the rDNA locus), and other chromosomes or loci as indicated. The average number of reads mapped on the chromosomes and *DUP4, DUP5, and DUPR* loci are presented. C. CNAtra output of read depth signals calculated from 3C-seq reads (black dots; bin size = 1 kb) and estimated copy numbers (red lines) of each chromosome in *C. tropicalis*. For regions whose estimated copy numbers are <1.5 or >2.5, their respective start and end coordinates (shown in black vertical text) as well as estimated copy numbers (shown in red horizontal text) are manually labelled. Black box represents a zoom in view of a region on ChrR (600-1000 kb) where a duplicated region shows an estimated copy number of ~4.

**Figure S5. Chromoblot, sequence coverage analysis, and haplotyping for validation of the chromosome-level genome assembly of *C. tropicalis.***

A. Schematic of the balanced heterozygous translocation between Chr1B and Chr4B. The *DUP4* locus is highlighted with the black striped box. The junction between Chr1 and Chr4 on Chr1B and Chr4B are marked with black and purple arrows, respectively. B. Contact probability heatmaps (bin size = 10 kb) of Chr1 and Chr4 of *C. tropicalis* showing a balanced translocation as evidenced by a butterfly-like pattern (chromatin contacts split into two blocks) in the interchromosomal area. The 3C-seq reads were mapped to Assembly2020 (top; original) with Chr1A and Chr4A genomic sequences (Fig. 1C). We have also mapped the 3C-seq reads to an alternate assembly (bottom) with Chr1B and Chr4B sequences. Alternate assembly has been generated by exchanging the genomic sequences at the translocation breakpoint in Chr1 and Chr4. Coordinate of translocation was mapped using two *de novo* assembled contigs supporting the junctions. Chromosome labels and their corresponding ideograms are shown on the heatmap. Colorbar represents the contact probability in log2 scale. E. An ethidium bromide stained gel image and phosphorimages obtained from Southern hybridization using a probe from part of Chr1, which is exchanged with Chr4 (probe F) and a second probe from part of Chr4, that is exchanged with Chr1 (Probe E) (Table S9). Black triangles point to the genomic coordinates of the probes used. C. IGV tracks showing 3C-seq (blue) and SMRT-seq coverage (yellow) across the translocation junctions on each of the unaltered homolog of Chr1 (black border) and Chr4 (purple border), respectively. D. An ethidium bromide stained gel image and phosphorimages obtained from

Southern hybridization using centromere-proximal probes from Chr1 (Probe C) and ChrR (Probe D). E. A synteny dot-plot comparing the colinearity between the chromosomes and the FALCON-generated contigs (labeled as a-l). Five very short contigs are denoted by an asterisk. The enlarged version of the dot plot for these contigs is shown on the right panel. The dot-plot was generated using Symap.

**Figure S6. Partial conservation of a LOH block in each of the *C. albicans, C. tropicalis* and *C. sojae* genome.**
The circos tracks represent the SNP density, positions of the centromeres, Indel density, Illumina sequence coverage (the sequence coverage at the rDNA loci is clipped for clearer representation and marked with an asterisk) as indicated. The ribbon plot was drawn by connecting the genomic coordinates of the conserved single copy orthologs between *C. tropicalis* and *C. sojae* (teal), and *C. tropicalis* and *C. albicans* (purple).

**Figure S7. Analysis of 3C-seq data reveals interchromosomal and intrachromosomal telomeric contacts in *C. tropicalis* genome.**
A. Histogram of all interchromosomal interactions (excluding zero values; gray) was plotted from the 3C-seq contact probability matrix (bin size = 2 kb) of *C. tropicalis*. The mean value of all interchromosomal interactions is indicated by the black vertical line. A cartoon of interchromosomal telomeric interactions (dotted curves) is depicted above the histogram as the interactions between telomeres of different chromosomes. The mean value of interchromosomal telomeric interactions is indicated by the blue vertical line in the histogram. The interchromosomal telomeric interactions are significantly greater than all interchromosomal interactions (Mann-Whitney U test P value = $1.129\times10^{-11}$). B. Histogram of all intrachromosomal long-range (>100 kb) interactions (excluding zero values; gray) was plotted from chromosome-wide contact probability matrices (bin size = 2 kb) of *C. tropicalis*. A cartoon of intrachromosomal telomeric interactions (dotted curves) is depicted above the histogram as the interactions between two telomeres of the same chromosome. *Inset*, a cartoon depicting long-range interactions (gray blocks) and intrachromosomal telomeric interactions (blue blocks) in a chromosome-wide matrix. A long-range interaction is defined as the *cis* interaction between two loci separated by a distance of >100 kb. Intrachromosomal telomeric interactions are computed as

*cis* interactions between loci whose distances to two telomeres have a sum of ≤10 kb. Note that the distances indicated in the cartoon for 100 kb and 10 kb are not drawn to scale. The mean values of all long-range and intrachromosomal telomeric interactions are indicated by black and blue vertical lines, respectively. The intrachromosomal telomeric interactions are significantly greater than all long-range interactions (Mann-Whitney U test P value = $7.374 \times 10^{-11}$).

**Figure S8. Genome-wide synteny analysis between *C. albicans* and *C. tropicalis* suggests evidence of inter-centromere translocations in the last common ancestor.**

A. Synteny maps of *C. tropicalis* chromosomes (the lowermost line of each panel, marked by filled black circles numbered from 1 to R), with respect to *C. albicans* chromosomes (lines above the *C. tropicalis* chromosomes), in the order of Chr1 to ChrR (top to bottom) for all panels. Centromeres, black triangles. The ORFs (represented as beads) are color-coded: inverted, red and non-inverted, green. The more conserved the reciprocal best hits (RBH) are, the darker are the shades of red/green color. B. Zoom in view of the synteny relationship between the centromere proximal ORFs of *C. tropicalis* Chr6 with *C. albicans* Chr7. C. The zoom in view of RBH ORFs proximal to the centromeres of *C. tropicalis* as indicated in the figure, where each centromere is located at an ICSB. D. A scaled representation of the color-coded orthoblocks (relative to *C. tropicalis* chromosomes) and ICSBs (white lines) in *C. albicans* (Methods). Orthoblocks are defined as stretches of the target genome (*C. albicans*) carrying more than two syntenic ORFs from the same chromosome of the reference genome (*C. tropicalis*). The centromeres are represented with black arrowheads. E. Circos plot showing the ICSBs (purple lines on the outer-most circle) on *C. tropicalis* chromosomes (marked with black filled circles). The centromere proximal ORFs (10 ORFs on both sides) present in *C. tropicalis* are connected to their homologs present on *C. albicans* chromosomes (marked by purple filled circles) by color-coded lines (based on their origin). The positions of centromeres are marked with black lines of the inner-most circle in each chromosome. The genomic locations in *C. albicans* chromosomes showing the convergence of ORFs from at least two centromere-proximal loci of *C. tropicalis* are marked with red (proximal to the *C. albicans* centromere) and purple (a non-centromere locus) triangles. Note that all centromeres of *C. albicans* are proximal to

ORFs, homologs of which are proximal to centromeres of *C. tropicalis.* F - G. Circos plots showing the convergence of centromere proximal ORFs of *C. tropicalis* chromosomes near the centromeres on *C. albicans* chromosomes 3 (CaChr3) and 7 (CaChr7), respectively. Chromosomes of *C. tropicalis* and *C. albicans* are marked with black and purple filled circles at the beginning of each chromosome, respectively.

**Figure S9. Identification of HIR-associated centromeres in the CUG-Ser1 clade.** A. Schematic of the method used for the identification of putative centromeres in *C. sojae*, *C. viswanathii,* and *C. parapsilosis.* Putative centromeric loci in these species were tested for gene synteny with *C. tropicalis* (for *C. sojae* and *C. viswanathii*), presence of IRs, and overlap with intergenic/ORF-free regions. B. Genome-wide synteny of conserved ortholog pairs between *C. sojae* and *C. tropicalis.* C. Genome-wide synteny between conserved ortholog pairs between *C. viswanathii* and *C. tropicalis*. The location of the centromere on each chromosome is marked with a black bar. Chromosome numbers are marked at the beginning of each chromosome or contigs in colored filled circles (a to g are tig00000002, tig00000008, tig00000017, tig00000038, tig00000050, tig00016100, tig00000001 and h to n are NW_020797881.1, NW_020797885.1, NW_020797858.1, NW_020797886.1, NW_020797884.1, NW_020797877.1 and NW_020797878.1). Contigs that are either <100 kb in length or do not carry putative centromeres (for *C. sojae*) or duplicated in the genome assembly (for *C. viswanathii*) were excluded from this analysis. The chromosomal coordinates of the ortholog pairs are connected using lines. D. and E. Circos plots similar to that of B and C, showing 10 ORFs on both sides of each centromere of *C. tropicalis* connected to the corresponding genomic loci carrying homologs in *C. sojae* and *C. viswanathii,* respectively.

**Figure S10. Inter-species conservation of centromere DNA sequences of closely related *Candida* species.**
A. Bean plots showing the distribution of the percent sequence identity among the centromeric left repeat (LR), the central core (CC), and right repeat (RR) elements in *C. tropicalis* (*Ct*), *C. sojae* (*Cs*), *C. viswanathii* (*Cv*), and *C. parapsilosis* (*Cp*). The extent of sequence identity between all possible pairs of CC, LR, RR, and LR-RR pairs were calculated in blastn analysis using Clustal Omega. The means are

depicted as a horizontal black bar on each of the bean pod. B. Bean plots showing distribution and means (horizontal black bar on each of the bean pods) of percent sequence identity values obtained from pairwise DNA sequence alignment of all possible combinations for each of seven random loci (Random), centromeric left repeats (LR), central cores (CC) and right repeats (RR) between species-pairs as indicated, using Clustal Omega. The significance of difference between percent sequence identity of centromere elements and random loci for all three species pairs were tested using the Mann-Whitney U test ($P < 0.05$) and the P value summary for each comparison is represented with asterisks. C. Heatmap plot across the contigs carrying the putative centromeres showing the extent of enrichment of IR-motifs in *C. sojae* (a-g, as described in Figure S9B) and *C. viswanathii* (h-m, as described in Figure S9B). The locations of the centromeres are pointed with red arrowheads. D. IGV track images showing the extent of enrichment of IR-motif on the putative centromeres of *C. sojae* and *C. viswanathii.* E. A representative figure showing a zoom in view of the IR-motif distribution on *C. tropicalis CEN1* DNA. The motifs on the Crick strand (red) and Watson strand (blue) are color coded. F. Heatmap showing the percent of IR-motifs present in converging and diverging orientation with respect to the central core region for each of the HIR associated putative centromeres present in *C. sojae, C. tropicalis,* and *C. viswanathii.* G. The average number of IR-motifs per 250 bp on the IRs is plotted in the *y*-axis as a function of the distance from the start of CC (*x*-axis) for *C. tropicalis* (red), *C. sojae* (yellow), and *C. viswanathii* (blue). Bean plots were generated using BoxplotR.

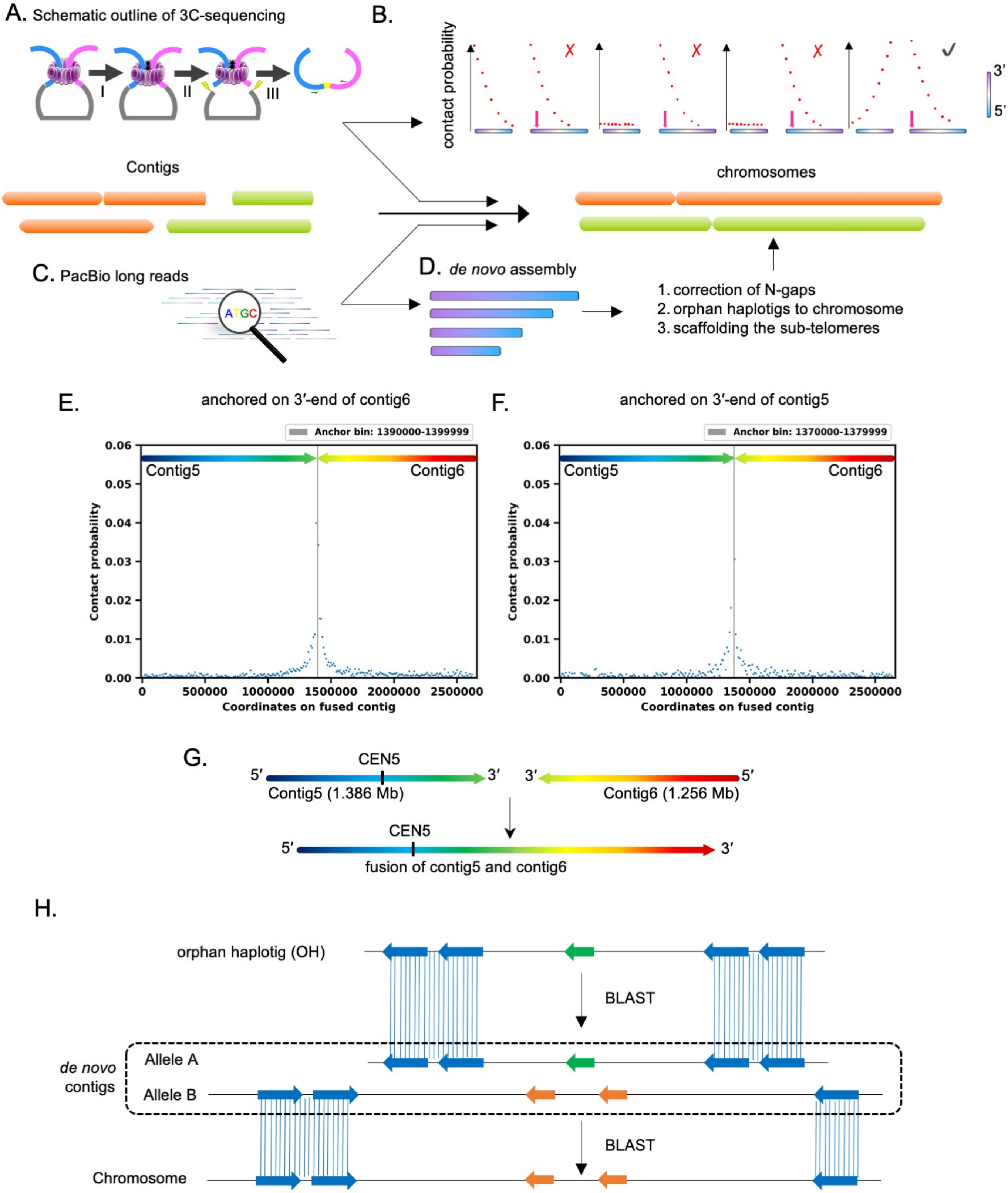

A. Schematic outline of 3C-sequencing
I
II
III
B.
contact probability
3'
5'
Contigs
chromosomes
C. PacBio long reads
ATGC
D. de novo assembly
1. correction of N-gaps
2. orphan haplotigs to chromosome
3. scaffolding the sub-telomeres
E.
anchored on 3′-end of contig6
Anchor bin: 1390000-1399999
Contig5
Contig6
Contact probability
Coordinates on fused contig
F.
anchored on 3′-end of contig5
Anchor bin: 1370000-1379999
Contig5
Contig6
Contact probability
Coordinates on fused contig
G.
CEN5
5′
3′
3′
5′
Contig5 (1.386 Mb)
Contig6 (1.256 Mb)
CEN5
5′
3′
fusion of contig5 and contig6
H.
orphan haplotig (OH)
BLAST
de novo contigs
Allele A
Allele B
BLAST
Chromosome


**Figure S1**

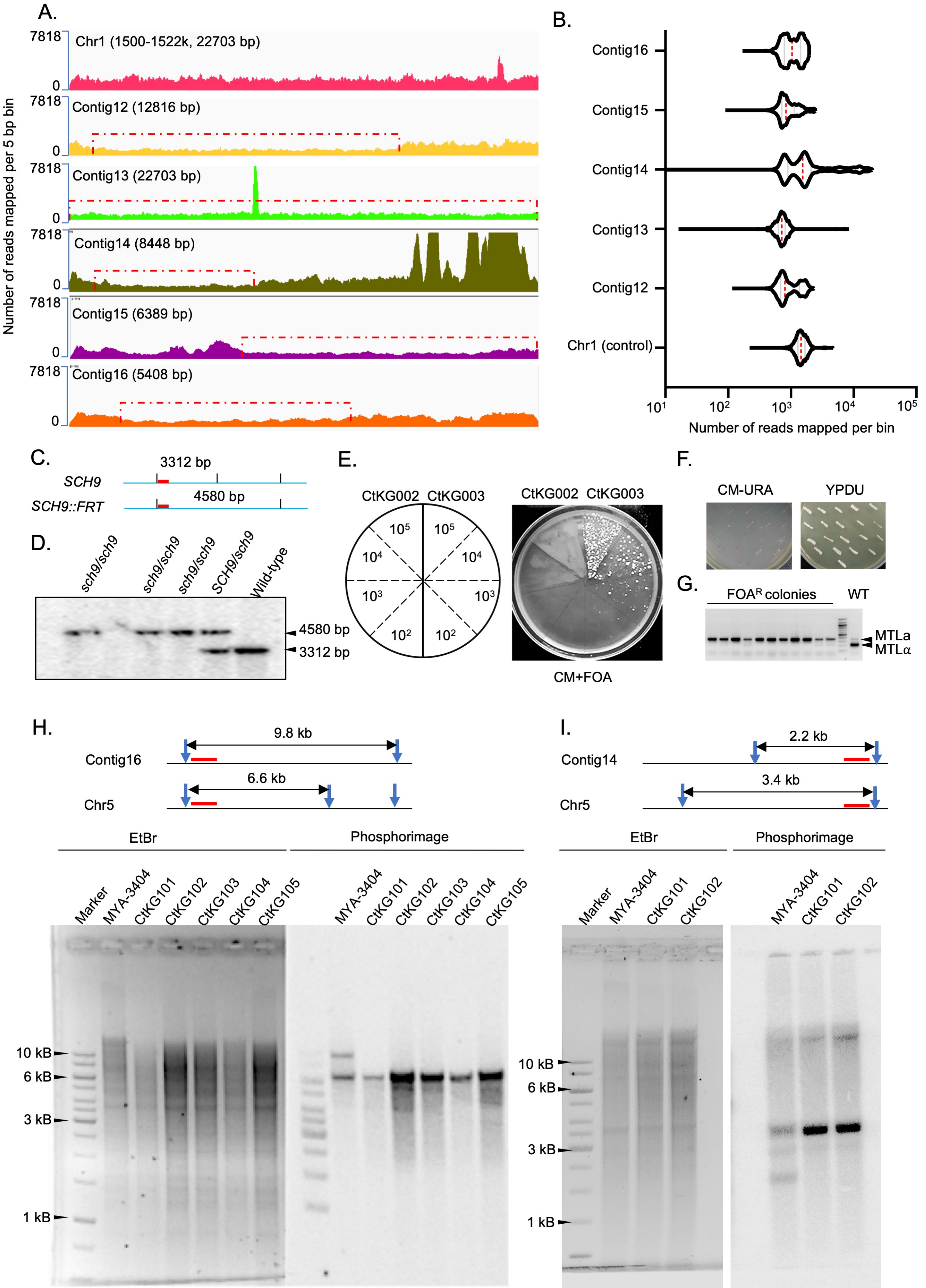

A.
Number of reads mapped per 5 bp bin
7818
0
Chr1 (1500-1522k, 22703 bp)
Contig12 (12816 bp)
Contig13 (22703 bp)
Contig14 (8448 bp)
Contig15 (6389 bp)
Contig16 (5408 bp)
B.
Contig16
Contig15
Contig14
Contig13
Contig12
Chr1 (control)
Number of reads mapped per bin
C.
SCH9
SCH9::FRT
3312 bp
4580 bp
D.
sch9/sch9
sch9/sch9
sch9/sch9
SCH9/sch9
Wild-type
4580 bp
3312 bp
E.
CtKG002
CtKG003
CM+FOA
F.
CM-URA
YPDU
G.
FOAR colonies
WT
MTLa
MTLα
H.
9.8 kb
Contig16
6.6 kb
Chr5
EtBr
Phosphorimage
Marker
MYA-3404
CtKG101
CtKG102
CtKG103
CtKG104
CtKG105
10 kB
6 kB
3 kB
1 kB
I.
2.2 kb
Contig14
3.4 kb
Chr5


**Figure S2**

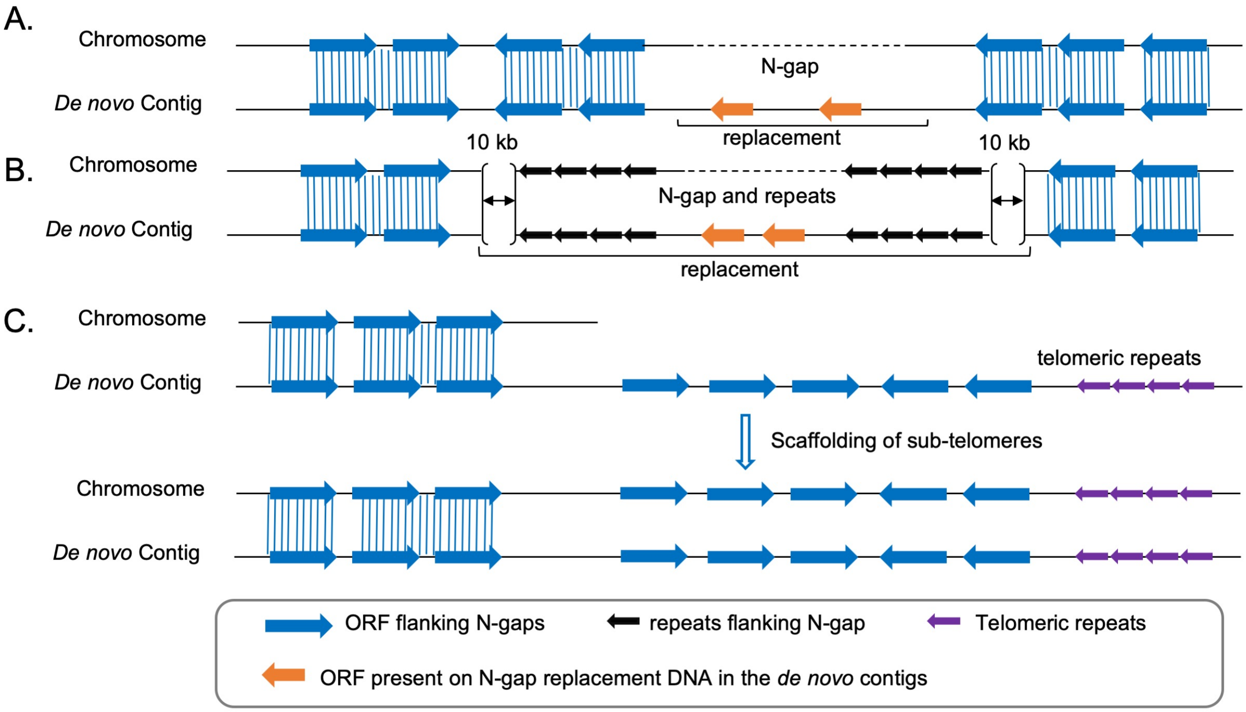
A.
Chromosome
De novo Contig
N-gap
replacement
B.
Chromosome
De novo Contig
10 kb
10 kb
N-gap and repeats
replacement
C.
Chromosome
De novo Contig
telomeric repeats
Scaffolding of sub-telomeres
Chromosome
De novo Contig
ORF flanking N-gaps
repeats flanking N-gap
Telomeric repeats
ORF present on N-gap replacement DNA in the de novo contigs

**Figure S3**

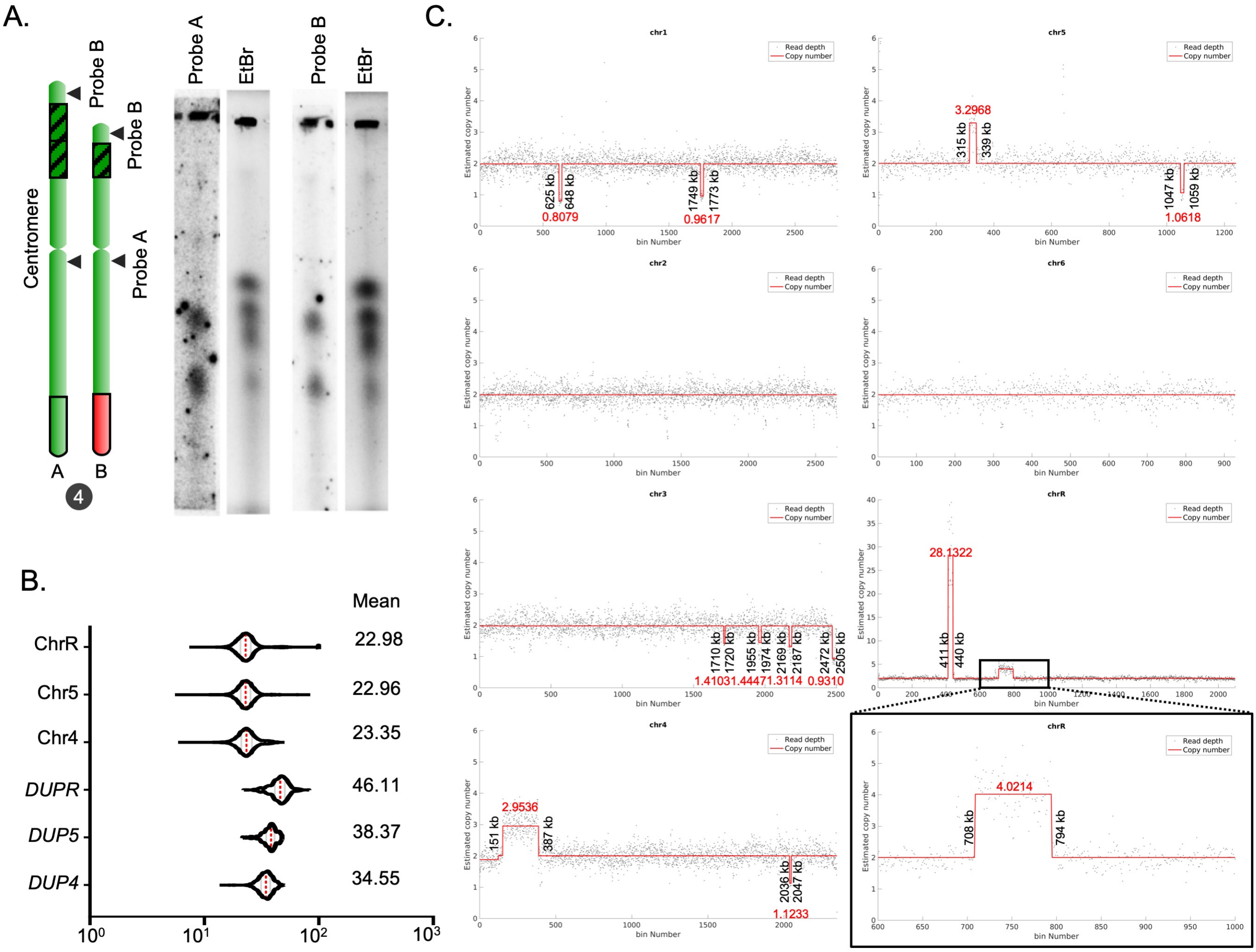
A.
Probe B
Probe B
Centromere
Probe A
A
B
4
Probe A
EtBr
Probe B
EtBr
B.
Mean
ChrR
22.98
Chr5
22.96
Chr4
23.35
DUPR
46.11
DUP5
38.37
DUP4
34.55
C.
chr1
625 kb
648 kb
0.8079
1749 kb
1773 kb
0.9617
chr5
3.2968
315 kb
339 kb
1047 kb
1059 kb
1.0618
chr2
chr6
chr3
1710 kb
1720 kb
1955 kb
1974 kb
2169 kb
2187 kb
2472 kb
2505 kb
1.41031.44471.3114 0.9310
chrR
28.1322
411 kb
440 kb
chr4
2.9536
151 kb
387 kb
2036 kb
2047 kb
1.1233
chrR
4.0214
708 kb
794 kb
Read depth
Copy number
Estimated copy number
bin Number

Figure S4

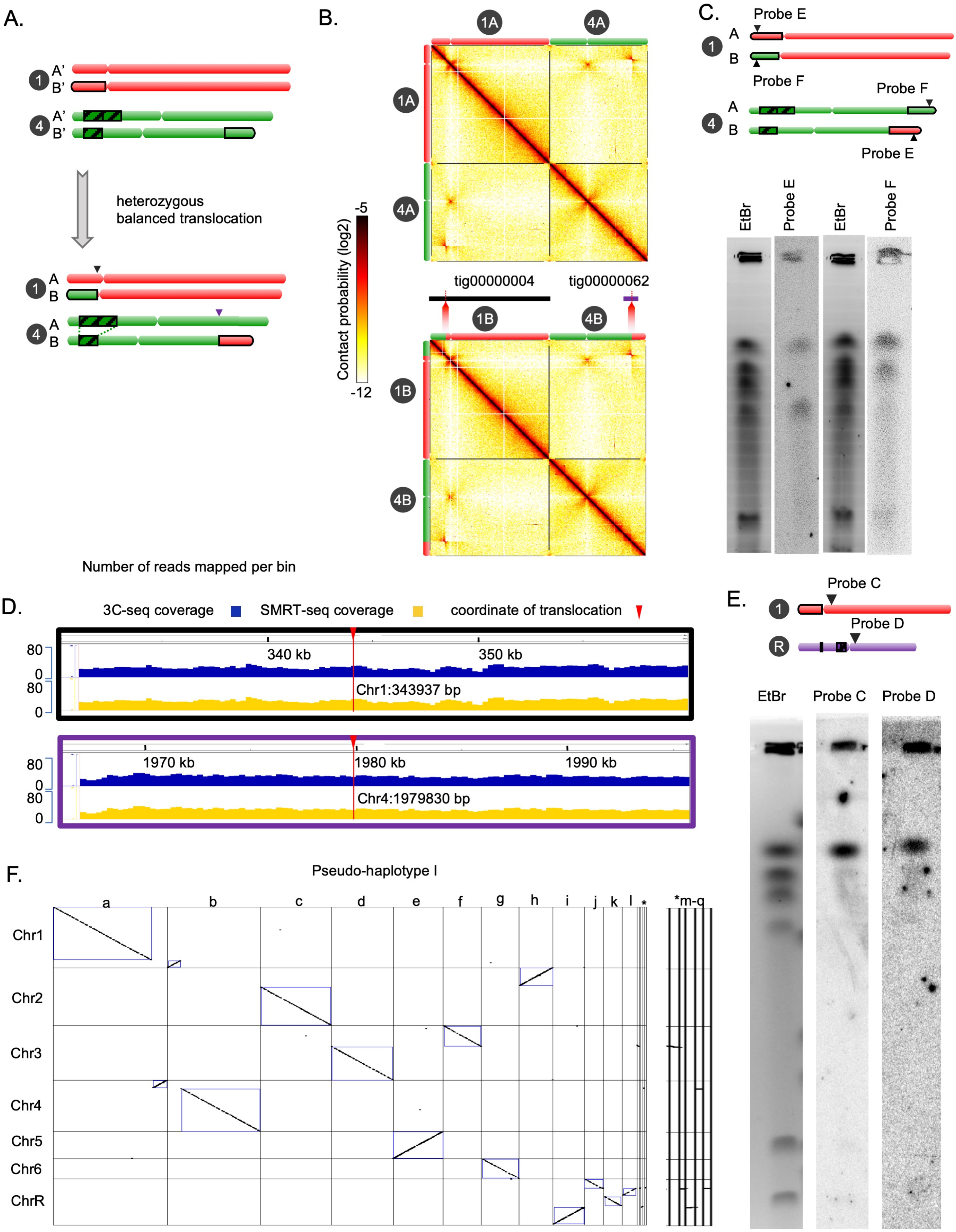
A.
A'
B'
A'
B'
heterozygous
balanced translocation
A
B
A
B
B.
1A
4A
1A
4A
Contact probability (log2)
-5
-12
tig00000004
tig00000062
1B
4B
1B
4B
C.
Probe E
A
B
Probe F
Probe F
A
B
Probe E
EtBr
Probe E
EtBr
Probe F
Number of reads mapped per bin
D.
3C-seq coverage
SMRT-seq coverage
coordinate of translocation
80
0
80
0
340 kb
350 kb
Chr1:343937 bp
80
0
80
0
1970 kb
1980 kb
1990 kb
Chr4:1979830 bp
E.
Probe C
Probe D
R
EtBr
Probe C
Probe D
F.
Pseudo-haplotype I
a
b
c
d
e
f
g
h
i
j
k
l
*
*m-q
Chr1
Chr2
Chr3
Chr4
Chr5
Chr6
ChrR

**Figure S5**

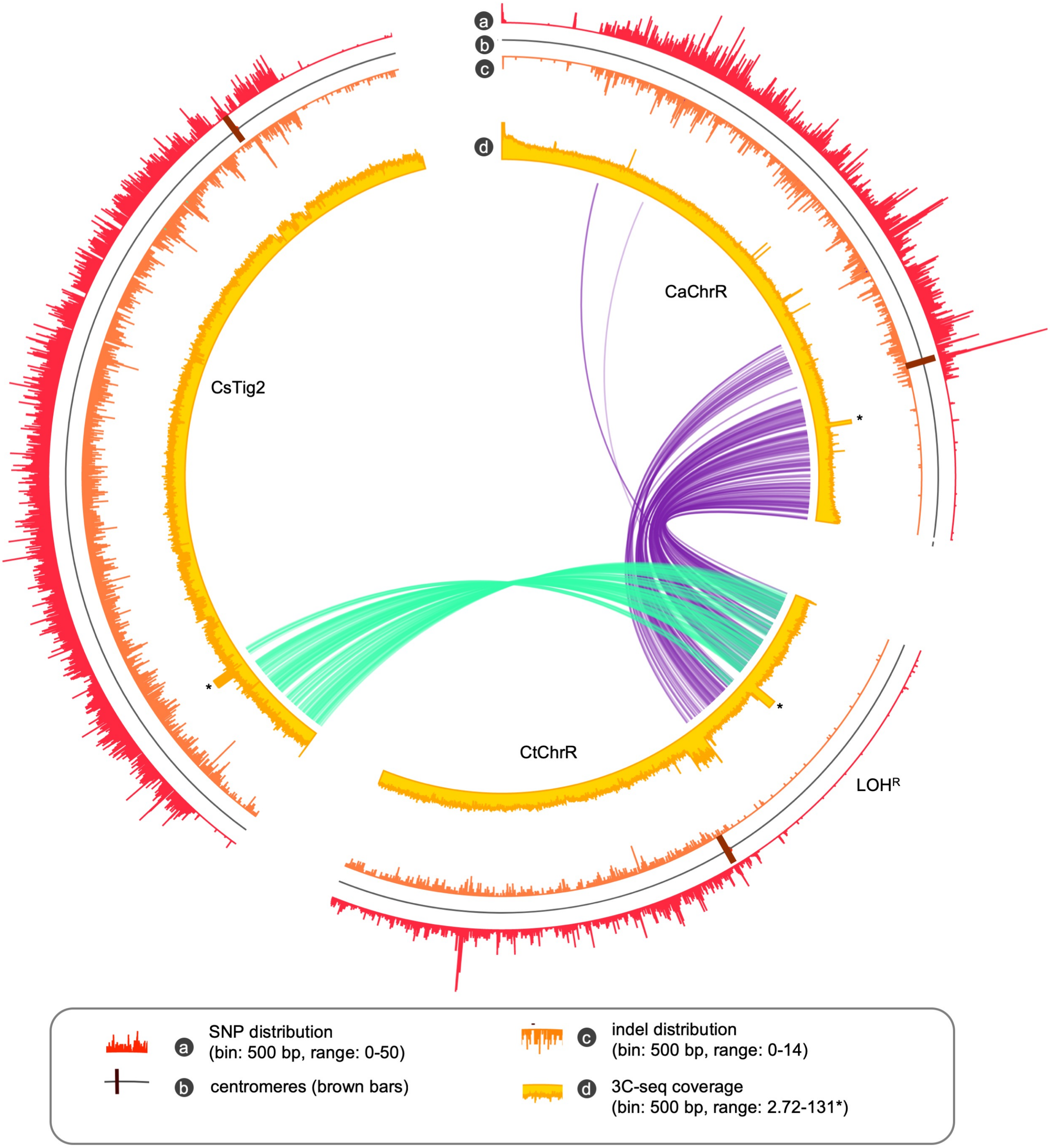
a
b
c
d
CaChrR
CsTig2
CtChrR
LOH$^{R}$
*
SNP distribution
(bin: 500 bp, range: 0-50)
centromeres (brown bars)
indel distribution
(bin: 500 bp, range: 0-14)
3C-seq coverage
(bin: 500 bp, range: 2.72-131*)

**Figure S6**

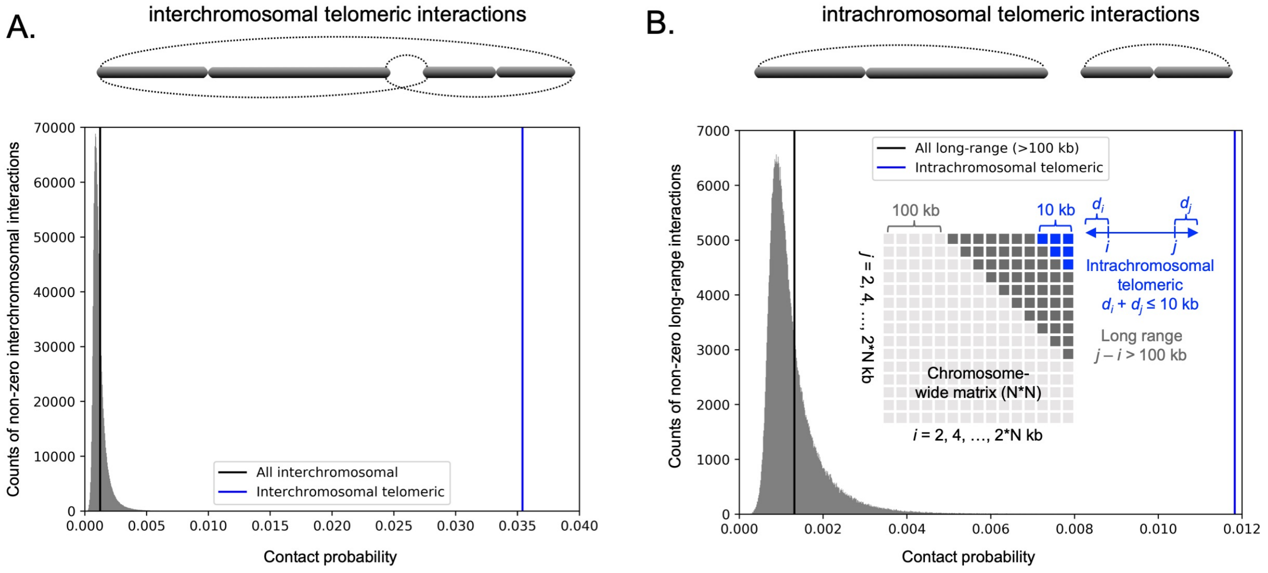
A.
interchromosomal telomeric interactions
Counts of non-zero interchromosomal interactions
Contact probability
All interchromosomal
Interchromosomal telomeric
B.
intrachromosomal telomeric interactions
Counts of non-zero long-range interactions
Contact probability
All long-range (>100 kb)
Intrachromosomal telomeric
100 kb
10 kb
Intrachromosomal telomeric
d_i + d_j ≤ 10 kb
Long range
j – i > 100 kb
Chromosome-wide matrix (N*N)
i = 2, 4, …, 2*N kb
j = 2, 4, …, 2*N kb

**Figure S7**

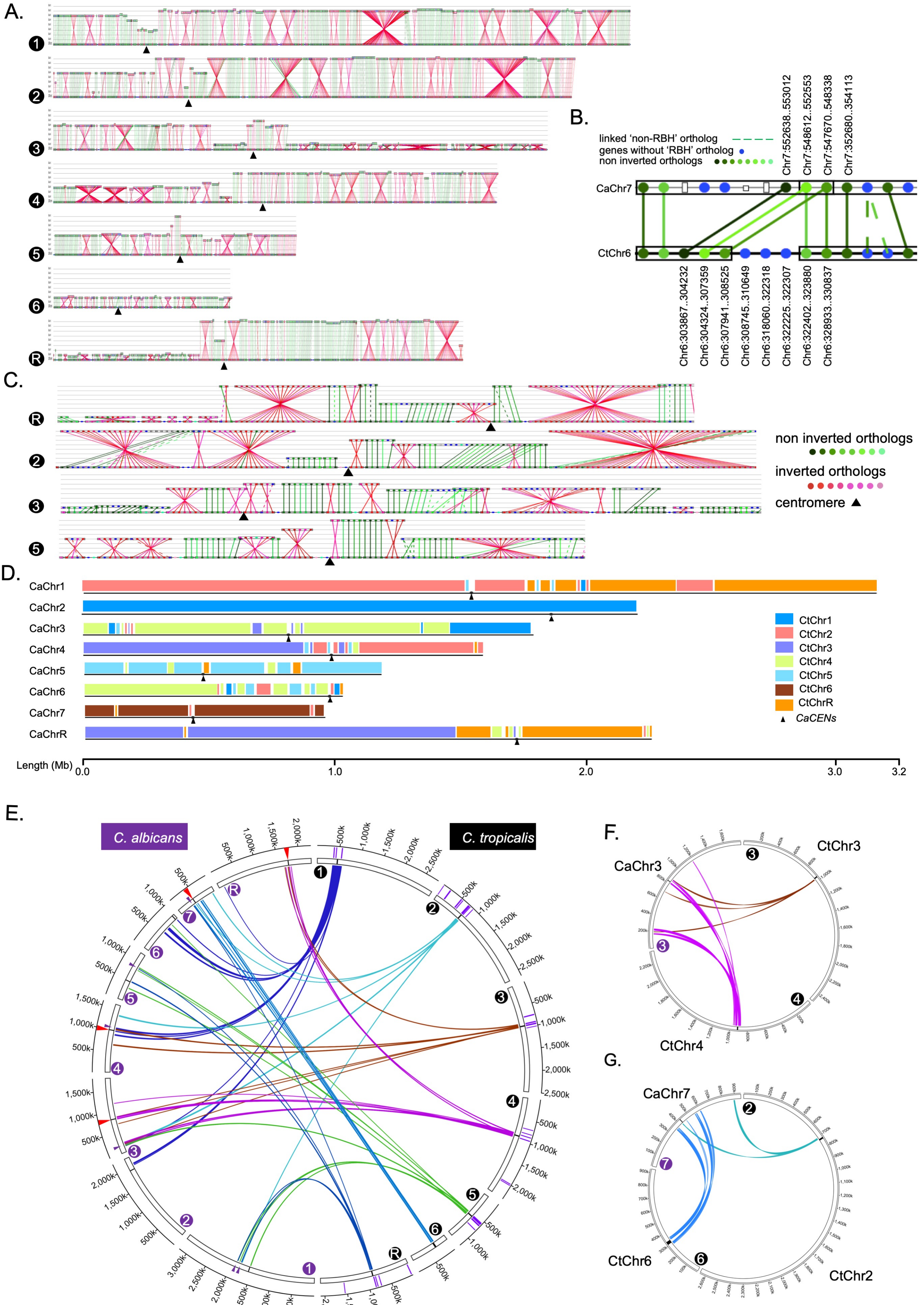
A.
B.
linked 'non-RBH' ortholog
genes without 'RBH' ortholog
non inverted orthologs
Chr7:552638..553012
Chr7:548612..552553
Chr7:547670..548338
Chr7:352680..354113
CaChr7
CtChr6
Chr6:303867..304232
Chr6:304324..307359
Chr6:307941..308525
Chr6:308745..310649
Chr6:318060..322318
Chr6:322225..322307
Chr6:322402..323880
Chr6:328933..330837
C.
non inverted orthologs
inverted orthologs
centromere
D.
CaChr1
CaChr2
CaChr3
CaChr4
CaChr5
CaChr6
CaChr7
CaChrR
CtChr1
CtChr2
CtChr3
CtChr4
CtChr5
CtChr6
CtChrR
CaCENs
Length (Mb)
0.0
1.0
2.0
3.0
3.2
E.
C. albicans
C. tropicalis
F.
CtChr3
CaChr3
CtChr4
G.
CaChr7
CtChr6
CtChr2

Figure S8

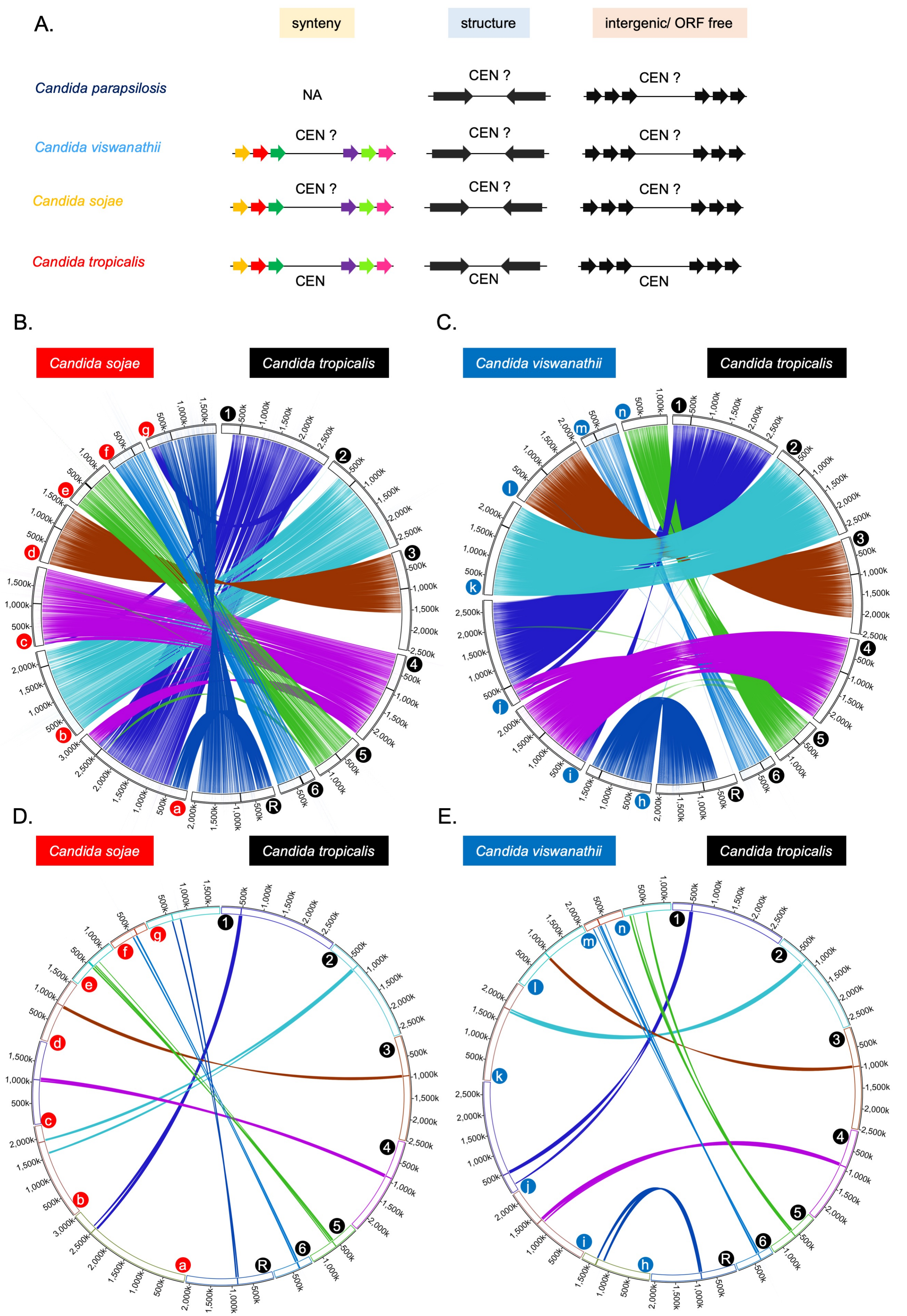
A.
synteny
structure
intergenic/ ORF free
Candida parapsilosis
NA
CEN ?
Candida viswanathii
Candida sojae
Candida tropicalis
CEN
B.
C.
D.
E.
Candida sojae
Candida tropicalis
Candida viswanathii

Figure S9

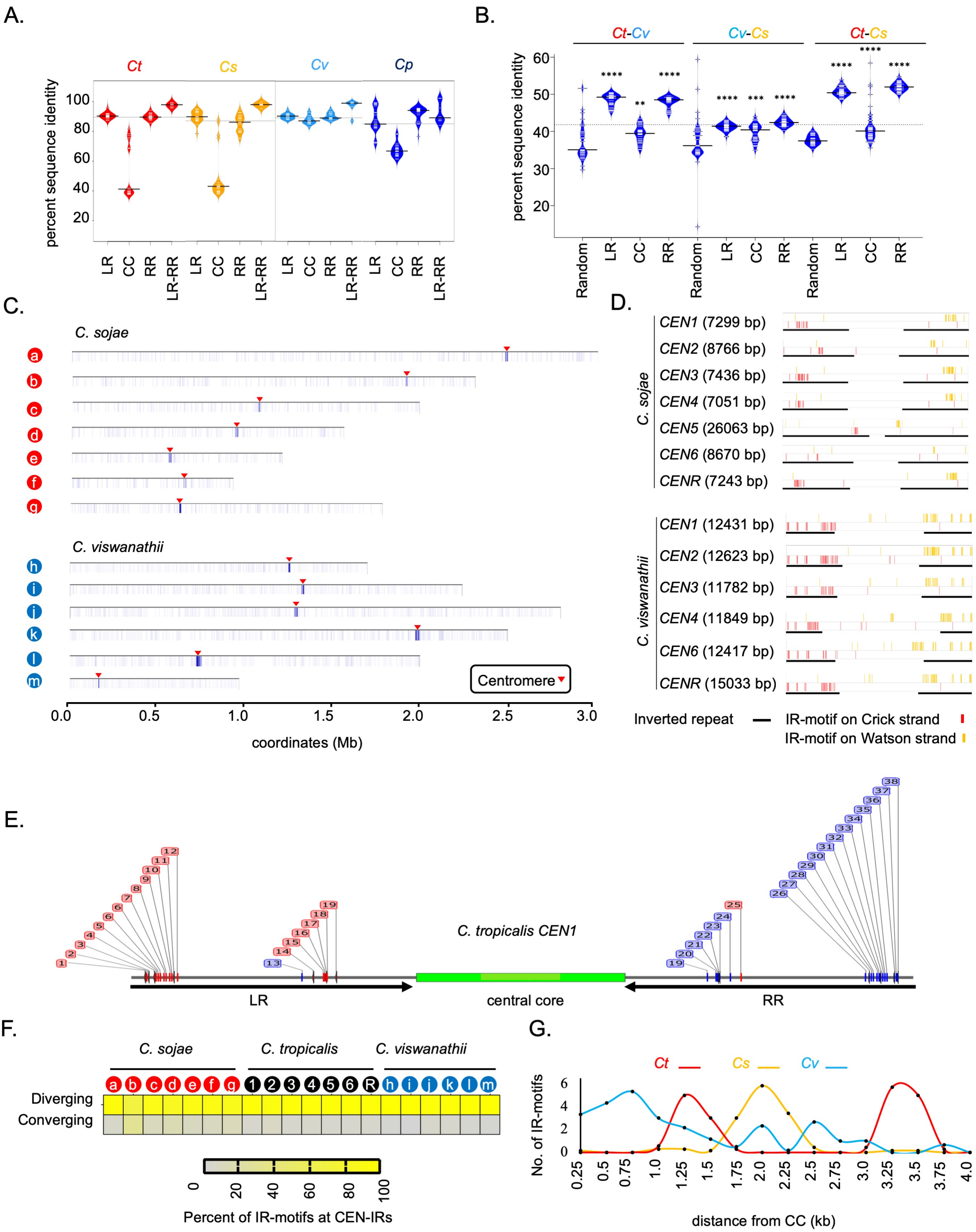
A.
Ct
Cs
Cv
Cp
percent sequence identity
100
80
60
40
20
LR
CC
RR
LR-RR
B.
Ct-Cv
Cv-Cs
Ct-Cs
percent sequence identity
60
50
40
30
20
Random
LR
CC
RR
C.
C. sojae
C. viswanathii
Centromere
0.0
0.5
1.0
1.5
2.0
2.5
3.0
coordinates (Mb)
D.
C. sojae
CEN1 (7299 bp)
CEN2 (8766 bp)
CEN3 (7436 bp)
CEN4 (7051 bp)
CEN5 (26063 bp)
CEN6 (8670 bp)
CENR (7243 bp)
C. viswanathii
CEN1 (12431 bp)
CEN2 (12623 bp)
CEN3 (11782 bp)
CEN4 (11849 bp)
CEN6 (12417 bp)
CENR (15033 bp)
Inverted repeat
IR-motif on Crick strand
IR-motif on Watson strand
E.
C. tropicalis CEN1
LR
central core
RR
F.
C. sojae
C. tropicalis
C. viswanathii
Diverging
Converging
0
20
40
60
80
100
Percent of IR-motifs at CEN-IRs
G.
Ct
Cs
Cv
No. of IR-motifs
distance from CC (kb)

**Figure S10**

Table S1: Assembly C with 12 contigs.

| Sl. No. | Contig No. | Length (bp) | CEN* | No. of N-gaps | Feature |
|---|---|---|---|---|---|
| 1 | 2 | 2797331 | 9 | 20 | NA |
| 2 | 1 | 2474493 | 1 | 15 | NA |
| 3 | 10 | 2342205 | 3 | 22 | NA |
| 4 | 11 | 2098823 | 4 | 15 | rDNA locus |
| 5 | 5 | 1379441 | 5 | 17 | Not observed in CHEF gel |
| 6 | 6 | 1255791 | None | | |
| 7 | 8 | 1225766 | 8 | 7 | MTLa |
| 8 | 7 | 921083 | 7 | 8 | NA |
| 9 | 12 | 12816 | None | - | MTLalpha |
| 10 | 14 | 8448 | None | - | NA |
| 11 | 15 | 6389 | None | - | NA |
| 12 | 16 | 5408 | None | - | NA |
| Total | 13 | 14527994 | 7 | 104 | |

*as reported in Chatterjee G. *et al.,* 2016.

Table S2: Assembly of sub-telomeres and filling up N-gaps in the genome assembly of *C. tropicalis* using *de novo* assembled contigs.

| Chr | 5′ tel | 3′ tel | Total N-gaps | Filled N-gap using strategy-I | Filled N-gaps using strategy-II |
|---|---|---|---|---|---|
| 1 | Tig4 | Tig28 | 20 | 14 | 6 |
| 2 | Tig224 | Tig10 | 17 | 14 | 3 |
| 3 | Tig236 | Tig2 | 15 | 11 | 4 |
| 4 | Tig4 | Tig238 5′ | 22 | 14 | 8 |
| 5 | Tig251 | Tig1909 | 7 | 7 | 0 |
| 6 | Tig110 5′ | Tig110 3′ | 8 | 4 | 4 |
| R | Tig254 | Tig244 | 15 | 14 | 1 |
| Total | | | 104 | 78 | 26 |

Table S3: Statistics for different versions of genome the assembly of *C. tropicalis* (MYA-3404) generated in this study

| Parameters | Assembly B | Assembly C | *De novo* (Canu) | *De novo* (FALCON) | Assembly2020 |
|---|---|---|---|---|---|
| asm_contigs | 16 | 12 | 135 | 17 | 7 |
| asm_esize | 1948827 | 2042279 | 1074242 | 1651545 | 2305488 |
| asm_max | 2797331 | 2797331 | 2853807 | 2860637 | 2835428 |
| asm_mean | 907756 | 1210666 | 138616 | 874119 | 2087075 |
| asm_median | 419327 | 1225766 | 35218 | 784244 | 2347188 |
| asm_min | 5408 | 5408 | 16806 | 30345 | 927416 |
| asm_n50 | 2216334 | 2342205 | 755598 | 1541447 | 2504192 |
| asm_n90 | 921083 | 1225766 | 35218 | 483039 | 1239682 |
| asm_n95 | 802573 | 921083 | 29731 | 464065 | 927416 |
| asm_total_bp | 14524098 | 14527994 | 18713202 | 14860026 | 14609527 |

Table S4: A comparative analysis of Assembly A and the improved Assembly2020 of *C. tropicalis*

| Features | Assembly A | Assembly2020 |
|---|---|---|
| Contigs/Chromosomes | 24 (23 and mtDNA) | 8 (7 and mtDNA) |
| Protein coding genes | 6254 | 6136 |
| tRNA genes | 187 | 190 |
| SNP density | 1 in 576 bp | 1 in 388 bp |
| Indel density | NA | 1 in 1340 bp |
| long heterozygous loci | 0 | 5 |
| Large CNVs | 0 | 3 |
| Telomeres | 0 | 14 |
| Total length | 14630139 bp | 14609527 bp |
| N50 | 1654078 bp | 2504192 bp |
| N90 | 498422 bp | 1239682 bp |
| Completeness (BUSCO) | 1255 out of 1315 | 1278 out of 1315 |
| N-gaps | 104 | 0 |

Table S5: Features of centromere DNA elements in *C. sojae**

| *CEN** | outer LR (bp) | left repeat (bp) | central core (bp) | right repeat (bp) | outer RR (bp) |
|---|---|---|---|---|---|
| *CENR* | 1389 | 2649 | 1967 | 2661 | 1386 |
| *CEN1* | 3957 | 2691 | 2138 | 2594 | 4009 |
| *CEN2* | ND | 3363 | 2111 | 3333 | ND |
| *CEN4* | ND | 2493 | 2068 | 2523 | ND |
| *CEN3* | ND | 2633 | 2207 | 2631 | ND |
| *CEN5*[#] | 12092 | | 2174 | 11797 | |
| *CEN6* | ND | 3313 | 2086 | 3312 | ND |

*syntenic to *CtCENs*; ND, Not detected; [#]Outer repeat is fused with the IR flanking the central core

Table S6: Features of centromere DNA elements in *C. viswanathii*

| *CEN** | left repeat (bp) | central core (bp) | right repeat (bp) |
|---|---|---|---|
| *CEN1* | 3238 | 5955 | 3238 |
| *CEN2* | 3792 | 5180 | 3651 |
| *CEN3* | 3228 | 5326 | 3228 |
| *CEN4* | 3115 | 5619 | 3115 |
| *CEN6* | 3165 | 6090 | 3162 |
| *CENR* | 2617 | 9798 | 2618 |

*syntenic to *CtCENs*

Table S7: Centromere coordinates used for identifying conserved DNA sequence motifs in *Candida* species

| Species | *CENs* | Coordinate |
|---|---|---|
| *C. albicans* | 1 | Ca22chr1A_C_albicans_SC5314:1561401-1569971 |
| | 2 | Ca22chr2A_C_albicans_SC5314:1923002-1930207 |
| | 3 | Ca22chr3A_C_albicans_SC5314:822280-826495 |
| | 4 | Ca22chr4A_C_albicans_SC5314:990403-998773 |
| | 5 | Ca22chr5A_C_albicans_SC5314:467755-473497 |
| | 6 | Ca22chr6A_C_albicans_SC5314:978304-983784 |
| | 7 | Ca22chr7A_C_albicans_SC5314:423402-429940 |
| | R | Ca22chrRA_C_albicans_SC5314:1741032-1748631 |
| *C. tropicalis* | 1 | Chr1:466,066-475,754 |
| | 2 | Chr2:725185-735300 |
| | 3 | Chr3:952377-962681 |
| | 4 | Chr4:906302-916031 |
| | 5 | Chr5:594006-604320 |
| | 6 | Chr6:308570-331012 |
| | R | ChrR:855697-865834 |
| *C. sojae* | 1 | Tig2:2493969-2501301 |
| | 2 | Tig8:1905358-1914164 |
| | 3 | Tig38:934048-941518 |
| | 4 | Tig17:1062429-1069512 |
| | 5 | Tig50:549697-575759 |
| | 6 | Tig16100:638309-647019 |
| | R | Tig1:623000-630276 |
| *C. viswanathii* | 1 | NW_020797858.1:473076-485506 |
| | 2 | NW_020797886.1:1829726-1842348 |
| | 3 | NW_020797884.1:937333-949114 |
| | 4 | NW_020797885.1:1322147-1333995 |
| | 6 | NW_020797877.1:349856-362272 |
| | R | NW_020797881.1:1345731-1360763 |

| Species | *CENs* | Coordinate |
|---|---|---|
| *C. parapsilosis* | 1 | HE605202:209,424-215,647 |
| | 2 | HE605203:362,588-368,324 |
| | 3 | HE605204:470,757-476,666 |
| | 4 | HE605205:1,281,284-1,287,683 |
| | 5 | HE605206:1,309,222-1,314,566 |
| | 6 | HE605207:658,126-664,775 |
| | 7 | HE605208:888,702-891,518 |
| | R | HE605209:470,949-477,776 |
| *C. dubliniensis* | 1 | Chr1:1,594,163-1,611,889 |
| | 2 | Chr2:1,942,441-1,947,217 |
| | 3 | Chr3:870,922-876,511 |
| | 4 | Chr4:1,028,245-1,036,391 |
| | 5 | Chr5:491,715-501,426 |
| | 6 | Chr6:1,001,458-1,009,565 |
| | 7 | Chr7:434,210-439,177 |
| | R | ChrR:1,713,450-1,722,609 |

Table S8: List of strains used in this study.

| Strain name | Genotype |
|---|---|
| SC5314 | *C. albicans* reference strain (Wild-type) |
| MYA3404 | *C. tropicalis* Clinical isolate (Wild-type) |
| CtKS102 | *ura3::FRT/ura3::FRT his1::FRT/his1::FRT arg4::FRT/arg4::FRT CSE4/CSE4::CSE4-TAP (CaHIS1)* |
| CtKG001 | *ura3::FRT/ura3::FRT his1::FRT/his1::FRT arg4::FRT/arg4::FRT CSE4/CSE4::CSE4-TAP (CaHIS1) sch9::FRT/sch9::FRT* |
| CtKG002 | *ura3::FRT/ura3::FRT his1::FRT/his1::FRT arg4::FRT/arg4::FRT CSE4/CSE4::CSE4-TAP (CaHIS1) sch9::FRT/sch9::FRT Chr5-497kb/ Chr5-497kb::CaURA3* |
| CtKG003 | *ura3::FRT/ura3::FRT his1::FRT/his1::FRT arg4::FRT/arg4::FRT CSE4/CSE4::CSE4-TAP (CaHIS1) Chr5-497kb/ Chr5-497kb::CaURA3* |
| CtKG101 | *ura3::FRT/ura3::FRT his1::FRT/his1::FRT arg4::FRT/arg4::FRT CSE4/CSE4::CSE4-TAP (CaHIS1) sch9::FRT/sch9::FRT* Chr5 monosomy Transformant 1 |
| CtKG102 | *ura3::FRT/ura3::FRT his1::FRT/his1::FRT arg4::FRT/arg4::FRT CSE4/CSE4::CSE4-TAP (CaHIS1) sch9::FRT/sch9::FRT* Chr5 monosomy Transformant 2 |
| CtKG103 | *ura3::FRT/ura3::FRT his1::FRT/his1::FRT arg4::FRT/arg4::FRT CSE4/CSE4::CSE4-TAP (CaHIS1) sch9::FRT/sch9::FRT* Chr5 monosomy Transformant 3 |
| CtKG104 | *ura3::FRT/ura3::FRT his1::FRT/his1::FRT arg4::FRT/arg4::FRT CSE4/CSE4::CSE4-TAP (CaHIS1) sch9::FRT/sch9::FRT* Chr5 monosomy Transformant 4 |
| CtKG105 | *ura3::FRT/ura3::FRT his1::FRT/his1::FRT arg4::FRT/arg4::FRT CSE4/CSE4::CSE4-TAP (CaHIS1) sch9::FRT/sch9::FRT* Chr5 monosomy Transformant 5 |
| NCYC-2606 | Environmental isolate of C. sojae (wild-type) |

Table S9: List of primers used in this study

| Primer name | Sequence | Purpose of use |
|---|---|---|
| Construction and confirmation of sch9/sch9 mutants | | |
| KG1 | ATGCGGTACCGTTGGTACCTTCTACAGATGC | Amplification and cloning of upstream homology region of SCH9 ORF |
| KG2 | AGTCCTCGAGAATGGGTGAGCAGATGATGG | |
| KG3 | ATGCCCGCGGGATGAAGAAATGCAACCAGCAG | Amplification and cloning of downstream homology region of SCH9 ORF |
| KG4 | ATGAGAGCTCCAAAATTGGAATCGTTAGAAACGG | |
| KG5 | CAACAATTTAACTTAACATGTGGCAC | PCR confirmation of Sch9::CaSAT1 |
| KG6 | TTCTGAAACTTGAAGGATTAGATAC | |
| KG7 | CAGTGGCTACAACTCAGAGCACGC | |
| KG8 | TTAGAGACACAAACGAACAATGTACC | |
| Construction and confirmation of URA3 integrated reporter strain | | |
| KG9 | GATCGGGCCCGGGGAACACCAACTTCAAAA | Amplification and cloning of upstream homology region of Chr5-497kb locus |
| KG10 | AGTCCTCGAGGAGAGTCATGACACACCACTTGTTG | |
| KG11 | ATCGCTGCAGGACTGGAACCTTATGTGAGGAGACAG | Amplification and cloning of downstream homology region of Chr5-497_kb locus |
| KG12 | AGCTGGATCCGACATCATGGATGAGCCTTGGTAG | |
| KG13 | GAGAAAAAGAAAGAGAAGGATTCTAAGG | PCR confirmation of Chr5-497_kb::URA3 transformants |
| KG14 | ATCCTTCTTCTTGGCCACCC | |
| KG15 | GGACTGGGAGGGTGCATTGG | Probe for Southern hybridization confirmation of the sch9/sch9 mutants |
| KG16 | CTATGTGGGCGTGTGATTGCGC | |
| KG17 | GATTTGGTATGAAAAGAGGAACTCTAAC | Amplification of MTLa locus |
| KG18 | CTACTAATTTTGAAACCATTTGGAGTCT | |
| KG19 | TAAAACATTAAGCATAGAGGACAAAGAA | Amplification of the MTLalpha locus |
| KG20 | AACTTCAAATGCAAAATGTAAAACATAC | |
| Probes for Southern hybridization experiments | | |
| KG21 | GTTATTGAAGAACCTAGAGGG | Contig14_Probe |
| KG22 | AGCTTGTAATTCAGGTGACA | |
| KG23 | CTACTCCACAGAAACAATCTCC | Contig16_Probe |
| KG24 | CAGGGATACACTTCTTATGACC | |
| KG25 | TGCAGTTGAAATCTCTTGGACC | Probe A |
| KG26 | GGATGTTGCGATCACTTTGG | |
| KG27 | GGAAAGATTCAGGATAAACCAATTG | Probe B |
| KG28 | CCATTTGACTTGCCCACTC | |
| KG29 | ATCATGAGAATACAAAGAGAAAGTT | Probe C |
| KG30 | AACTTTTGTAGTCTATCCAACTCTG | |
| KG31 | GTGCATACGTCACAGTTTGG | Probe D |
| KG32 | GATGCTATCATCTCAAACCAAGG | |
| KG33 | GGTGGTTACGGTACCAGATTG | Probe E |
| KG34 | CCGGCATTGATTCTGTTACC | |
| KG35 | GTTGTTATCCAGTGTACCAGTTG | Probe F |
| KG36 | GGGATTTCTGGTGGTTCAAC | |

Table S10: List of plasmids used in this study

| Name | Vector/ backbone | Modification | Length | References |
|---|---|---|---|---|
| pKG1 | pSFS2a | Upstream and downstream homology region of *SCH9 ORF is* cloned. | 8542 bp | This study |
| pKG2 | pBSCaURA3 | Upstream and downstream homology region of intergenic locus (Chr5_497_kb) is cloned. | 5039 bp | This study |
